# Couples in the deep: dissolved organic and microbial communities in the oxygenated hypolimnion of a deep freshwater lake

**DOI:** 10.1101/2025.09.23.678040

**Authors:** Morimaru Kida, Ayuri Ohira, Yusuke Okazaki, Yasuhiko T. Yamaguchi, Akiko S. Goto, Kazuhide Hayakawa, Hiroshi Nishimura

## Abstract

The interaction between dissolved organic matter (DOM) and microbial communities is a critical yet understudied driver of biogeochemical cycling in aquatic ecosystems. Understanding these interactions is essential for elucidating the chemical and microbial dynamics that sustain ecosystem functionings. Here, we combined non-target ultra high-resolution mass spectrometry-based environmental metabolome analysis with microbiome analysis to conduct the first comprehensive investigation of DOM-microbe linkages in both the epilimnion and oxygenated hypolimnion of a deep freshwater lake throughout the stratification period, with Lake Biwa (Japan) as a model system. To facilitate interpretation of DOM-microbe networks, we developed an integrated compound category classification (IC3) framework for assigning molecular formulae (MFs) to specific compound categories. Using a compositional data analysis framework, we identified specific MFs and bacterial taxa that covaried in the hypolimnion, which exhibited substantially more complex networks than the epilimnion. These networks encompassed 1705 out of the 1755 common MFs, representing the majority of total peak intensities, underscoring stronger DOM-microbe coupling in deep waters. Hypolimnion specialist bacteria were associated with specific MFs and co-ocuuring taxa, providing environmental metabolomic evidence for substrate preference and potential symbiotic relations. Notably, more than three-fourths of these MFs in relative abundance were classified as recalcitrant, including lipid-, lignin-, tannin-like, and carboxyl-rich alicyclic molecules, suggesting the capability of hypolimnion specialists to use or produce these compounds. Our study offers the first high-resolution insights into DOM-microbe associations in a deep freshwater lake and establishes a framework for more efficient and robust analyses of such interactions.

## 1. Introduction

Water bodies such as the oceans and lakes store vast amounts of carbon as organic compounds, collectively termed dissolved organic matter (DOM). Its total amount is more than the global biomass carbon and is comparable to the amount of atmospheric carbon dioxide (1). Therefore, the global carbon cycle and heat budget are considerably affected by the decomposition of only a small fraction of the DOM pool (2). This huge DOM pool contains not only C but also N, S, P, and other essential trace elements (e.g., Fe), thus the production and decomposition of DOM will have a significant impact on the dynamics of these trace elements and the ecosystem. Global warming is expected to increase water temperature, decrease dissolved oxygen concentration, and strengthen water stratification (3), all of which will affect DOM dynamics and the fate of microbes associated with it. Therefore, it is essential to elucidate the factors influencing DOM dynamics to predict the future state of material cycles and ecosystems in the aquatic environment and to take necessary countermeasures.

The interplay between DOM and microbes is a critical but grossly understudied component in material cycling in the aquatic environment. Aquatic biota release DOM spontaneously or through cell lysis (4). Water bodies also receive terrestrial OM from surrounding catchments which is typically enriched in aromatic material and lignin content. DOM exhibits molecular fingerprints reflecting all processes that it has received and can be considered as “environmental metabolome”. Microbes are known to be capable of utilizing various organic compounds in DOM for anabolism or catabolism (5). These microbial activities influence concentrations of organic compounds through consumption and production. At the same time, organic compounds can influence microbial abundance and composition as substrates or antibiotics (6). Mechanical understanding of this DOM-microbe interplay is essential in understanding the dynamics of the chemical and microbial sides and potentially ecosystem functioning that can arise as a result of these associations (7). Advances in analytical techniques have now enabled the detection of hundreds to thousands of molecular formulae (MFs) and microbial amplicon sequence variants (ASVs) in one water sample, especially using Fourier-transform ion cyclotron resonance mass spectrometry (FT-ICR MS) and 16S rRNA gene amplicon sequencing. By combining these techniques which can be considered in a unified framework of the “ecology of molecules” (8), the covariation between DOM molecules and microbes in water was studied in the last decade (7, 9–11). However, very few studies have conducted temporal analysis of this kind in the aquatic environment. To our knowledge, there are only four studies that investigated temporal dynamics of DOM-microbe interactions in natural aquatic systems with a similar resolution as the combination of FT-ICR MS and 16S rRNA gene amplicon sequencing (12–15). Three of them were conducted at fixed points in an estuary (monthly to twice-a-year sampling for all approximately 1.5 years) and the rest was in the North Sea (daily sampling for 20 days). There is no such detailed study in deep freshwater lakes or the oceans that covers both the sunlit epilimnion and dark hypolimnion. While a few previous studies have examined DOM-microbial interactions in multiple lakes, they were limited to surface water sampling at a single time point (11, 16). Furthermore, the data structure of mass spectrometry and sequencing data requires the use of “compositional data analysis” (17, 18) for valid association analysis of ASV-MF networks, but only a handful have adequately applied this methodology in studying their associations (19, 20).

The hypolimnion of lakes consists of a separated water layer under the thermocline during lake water stratification. In deep freshwater holomictic lakes with oligo- to mesotrophic conditions, the hypolimnion can remain oxygenated because heterotrophic oxygen demand does not exceed the stock of hypolimnetic oxygen (21). This oxygenated hypolimnion constitutes a large portion of the lake water volume where the mineralization and bacterial reworking of OM derived from the epilimnion occur, making it an important component of the lacustrine biogeochemical cycle. For instance, a significant contribution of epilimnetic DOM to the total hypolimnetic mineralization has been reported (21). Moreover, the hypolimnion of lakes houses unique microbial communities that are distinct from the epilimnion (22–25). These microbial communities are likely adapted to environments specific to the hypolimnion, such as a lack of solar irradiance, stabler water temperature, or lower oxygen concentration. Alongside these environmental conditions, influx (abundance and composition) of material derived from photosynthetic production in the epilimnion should also greatly influence the composition of hypolimnetic microbial assemblages that rely on it.. While substrate preferences of hypolimnion-dominant bacteria have been investigated (25, 26), related knowledge remains extremely scarce.

This study aimed to contribute to filling thes gaps (spatiotemporal dynamics of DOM-microbe interactions and substrate preferences of hypolimnion-dominant bacteria in lakes) by combining monthly sampling in the epilimnion and hypolimnion in a deep freshwater lake (Lake Biwa, Japan), environmental metabolome analysis by FT-ICR MS, and microbiome analysis by 16S rRNA gene amplicon sequencing, supplemented with a wide range of water chemistry parameters. Lake Biwa is a monomictic lake with an oxygenated hypolimnion that harbors one of the best-studied freshwater chemical and microbial ecosystems (21–23, 27–30). Thottathil et al. (2013) showed that the dynamics and processes of microbial-derived DOM in Lake Biwa were similar to those of the ocean, with humic-like DOM produced in the hypolimnion consuming oxygen (31), suggesting the potential of using the hypolimnion of a deep freshwater lake such as Lake Biwa as an analogue of the deep ocean. The hypolimnion of Lake Biwa during stratification can be regarded as a natural large-scale incubation experiment due to its stabilized conditions and the restricted material flux from the epilimnion caused by strong thermal and density stratification. By analyzing the spatiotemporal data with an explicit framework of compositional data analysis, we identified specific MFs and ASVs that covaried in the hypolimnion of Lake Biwa during stratification. This study also developed an integrated compound category classification, termed IC3, that is tailored for assignments of compound categories to MFs of DOM to aid the interpretation of ASV-MF networks.

## 2. Results

### 2.1. Basic water chemistry and SPE-DOM

The *in-situ* water profiling indicated that Lake Biwa underwent water stratification during May– December 2022 (Fig. 1d and 1e). Thus, our sampling covered the entire period of stratification. Water temperature showed clear seasonality in the epilimnion, while that in the hypolimnion remained constant at 7.5–8 °C (Fig. 1e). Dissolved oxygen concentrations were lowest at 85 m and kept decreasing during stratification (but > 2 mg L^-1^) (Fig. 1d), suggesting remineralization of organic matter at the deep. Dissolved organic carbon (DOC) and particulate organic carbon (POC) concentrations were higher in the epilimnion compared to the hypolimnion, with DOC peaks in late spring (May) and late summer (October), indicating photosynthetic production and accumulation of DOM during stratification (Fig. 1ab). The prokaryotic cell abundance was 0.85 to 4.72 (average = 1.87) 10^6^ cells mL^-1^ and higher in the epilimnion as in DOC and POC (Fig. 1c). The cell abundance was highest in summer (July–August) (Fig. 1c).

**Fig. 1.**
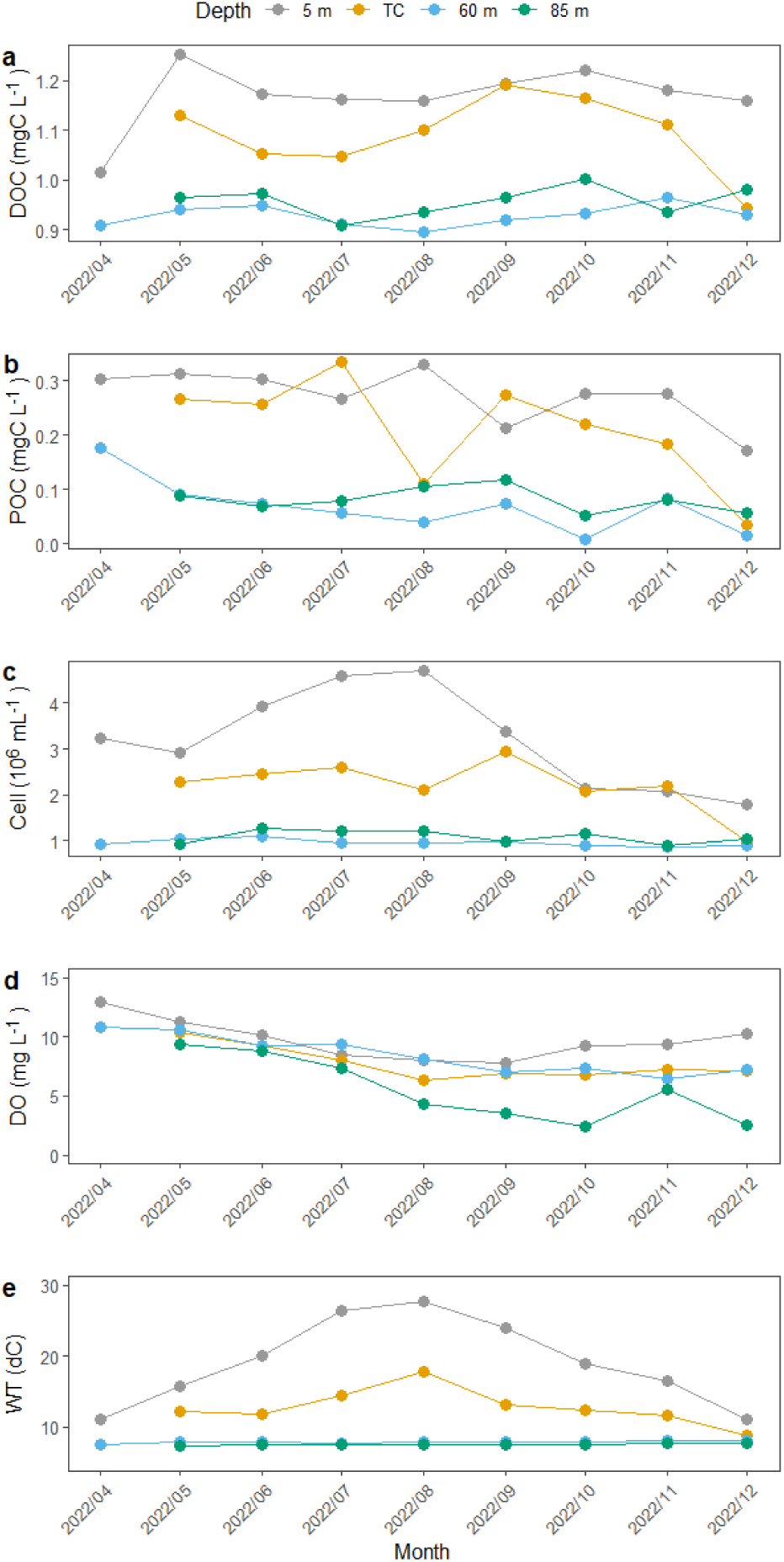
Dissolved organic carbon (a), particulate organic carbon (b), cell abundance (c), dissolved oxygen (d), and water temperature (e) data by different depths. In April of 2022, samples were not collected at the thermocline (TC) and 85 m.

Out of 6886 unique molecular formulae (MFs) detected by FT-ICR MS, 1755 MFs were common to all samples (2983–4004 MFs detected per sample), representing 88% of total peak intensities. FT-ICR MS broadband spectra of all samples were very similar and typical of natural DOM, with a bell-shaped distribution of mass peaks between approximately 150 m/z and 600 m/z. The extraction efficiency of DOC was not correlated with any environmental parameters. The rank abundance curve of solid phase-extracted DOM (SPE-DOM) was very steep, indicating that a small percentage of MFs was dominating overall chemical community composition (Fig. 2ac). There were little differences in species accumulation curves between different depths, although slightly more MFs were detected in the epilimnion (Fig. 2e). SPE-DOM chemical diversity quantified as ENS varied almost two-fold (380–664) and showed a clear unimodal, seasonal pattern, being highest in late summer (September), while differences in diversity between depths were small (Fig. 3a). Consistent trends in increasing molecular weight (m/z), aromaticity (modified aromaticity index, AI_mod_), and unsaturation (double bond equivalent, DBE and decreasing H/C) of SPE-DOM were observed with increasing depths, but these differences were overall small (Table 1). There were little differences in the elemental composition of SPE-DOM between depths (Table 1). Lake Biwa SPE-DOM was enriched in Lipid MFs, followed by carboxyl-rich alicyclic molecules (CRAM), Lignin, Tannin, Aromatic, Peptide, and small contributions from Carbohydrate and Amino sugar MFs (Table 1). Similarly, CHO-only MFs were dominant, followed by CHON-, CHOS-, CHONS-, CHOP-, CHOSP, and CHONP-containing MFs (Table 1). Regarding compound categories, %CRAM clearly increased while %Lipid decreased with depth (Table 1). Low aromaticity and relatively high nitrogen (N) content of SPE-DOM reflect a predominantly autochthonous origin of DOM in Lake Biwa (Table 1).

**Fig. 2.**
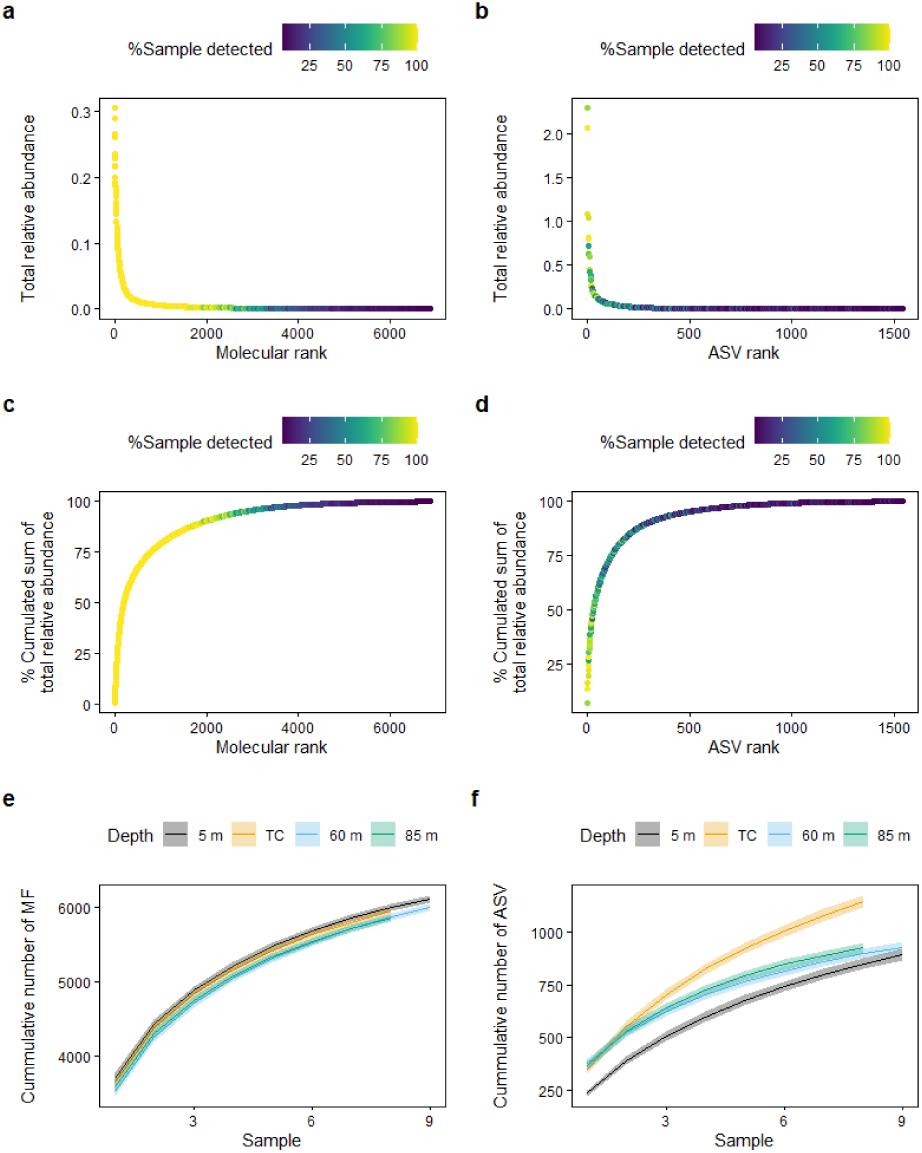
Rank abundance curves (a–d) and species accumulation curves (e and f) for solid-phase extracted dissolved organic matter (SPE-DOM; left panels) and 16s rRNA-based microbiome (right panels). In (a–d), coloring indicates the percentages of samples in which each molecular formula or amplicon sequence variant (ASV) was found across the dataset, while “total relative abundance” indicates the sum of sample-wise relative abundance across the dataset. In (e and f), the shaded areas represent 95% confidence intervals for each depth.

**Fig. 3.**
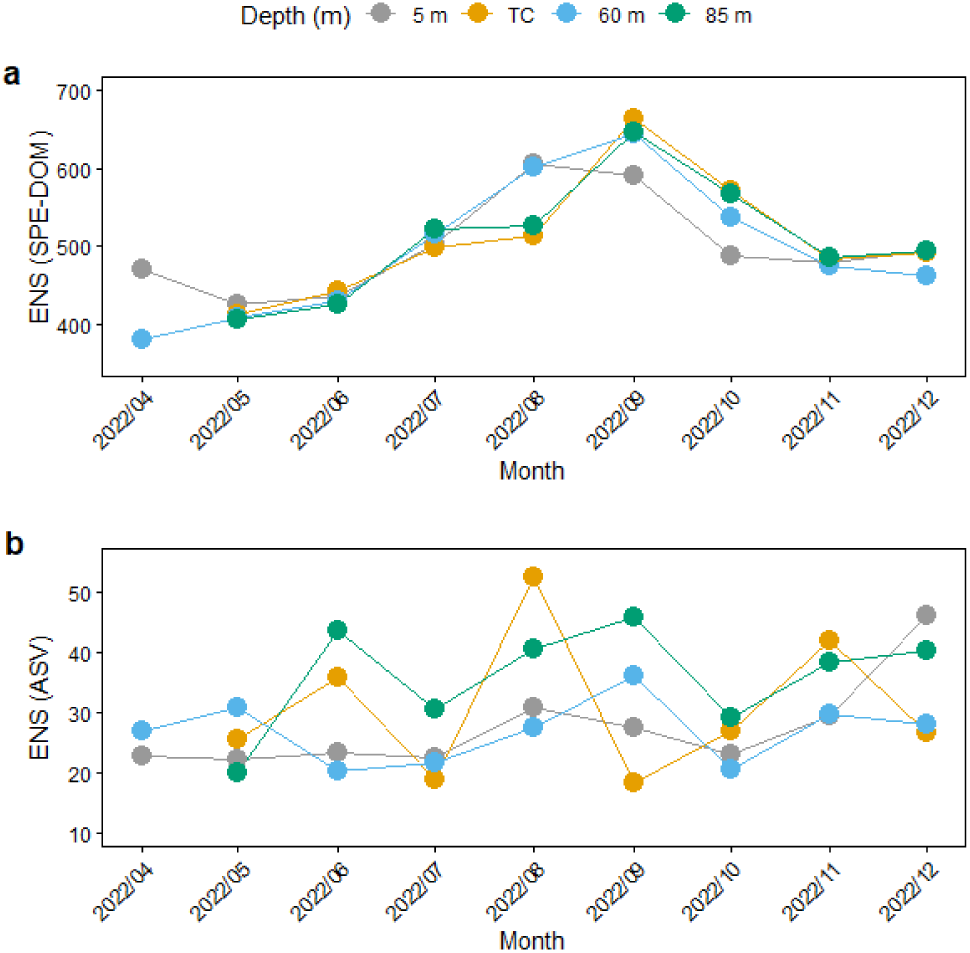
Diversity of SPE-DOM (a) and microbiome (b) quantified as the effective number of species by different depths.

**Table 1.** Summary of the intensity-weighted average (w.a.) of molecular properties and relative abundance (%RA) of compound categories and elemental classes.

|  | 5 m | TC | 60 m | 85 m |
| --- | --- | --- | --- | --- |
| H/C (w.a.) | 1.29 | 1.28 | 1.27 | 1.27 |
| O/C (w.a.) | 0.465 | 0.471 | 0.469 | 0.474 |
| m/z (w.a.) | 356.8 | 360.3 | 362.6 | 363.5 |
| AI<sub>mod</sub> (w.a.) | 0.232 | 0.237 | 0.241 | 0.242 |
| DBE (w.a.) | 7.18 | 7.34 | 7.44 | 7.49 |
| Amino sugar (%RA) | 0.23 | 0.21 | 0.21 | 0.21 |
| Aromatic (%RA) | 3.70 | 3.84 | 3.75 | 4.00 |
| CRAM (%RA) | 29.64 | 31.97 | 32.64 | 33.65 |
| Carbohydrate (%RA) | 0.93 | 0.85 | 0.72 | 0.75 |
| Lignin (%RA) | 13.94 | 13.90 | 14.97 | 14.36 |
| Lipid (%RA) | 44.06 | 41.33 | 39.83 | 38.75 |
| Peptide (%RA) | 3.33 | 3.26 | 3.19 | 3.28 |
| Tannin (%RA) | 4.18 | 4.64 | 4.69 | 5.00 |
| CHO (%RA) | 78.66 | 78.35 | 78.84 | 78.21 |
| CHON (%RA) | 16.53 | 16.64 | 16.29 | 16.73 |
| CHOS (%RA) | 3.68 | 3.83 | 3.76 | 3.89 |
| CHOP (%RA) | 0.29 | 0.28 | 0.26 | 0.29 |
| CHONS (%RA) | 0.47 | 0.48 | 0.43 | 0.45 |
| CHONP (%RA) | 0.16 | 0.18 | 0.20 | 0.20 |
| CHOSP (%RA) | 0.21 | 0.25 | 0.22 | 0.23 |

A clear seasonal pattern in the SPE-DOM composition was observed. The first two principal components of principal component analysis (PCA) explained 38% of the total variance in FT-ICR MS data (Fig. 4a). Seasonal samples were distributed in the PCA 2D ordination space counterclockwise from April to December (Fig. 4a). These sample distributions were correlated with molecular properties such as elemental ratios (O/C, H/C), relative abundance of compound categories, indices related to aromaticity and unsaturation (SUVA_254_, AI_mod_ and DBE), and chemical diversity of SPE-DOM assessed as effective number of species (ENS) (Fig. 4a). As environmental variables, dissolved oxygen (DO), pH, and chlorophyll fluorescence were significantly correlated (Fig. 4a). Consistent with the result in Fig. 3, ENS were higher in samples collected in summer (Fig. 4a).

**Fig. 4.**
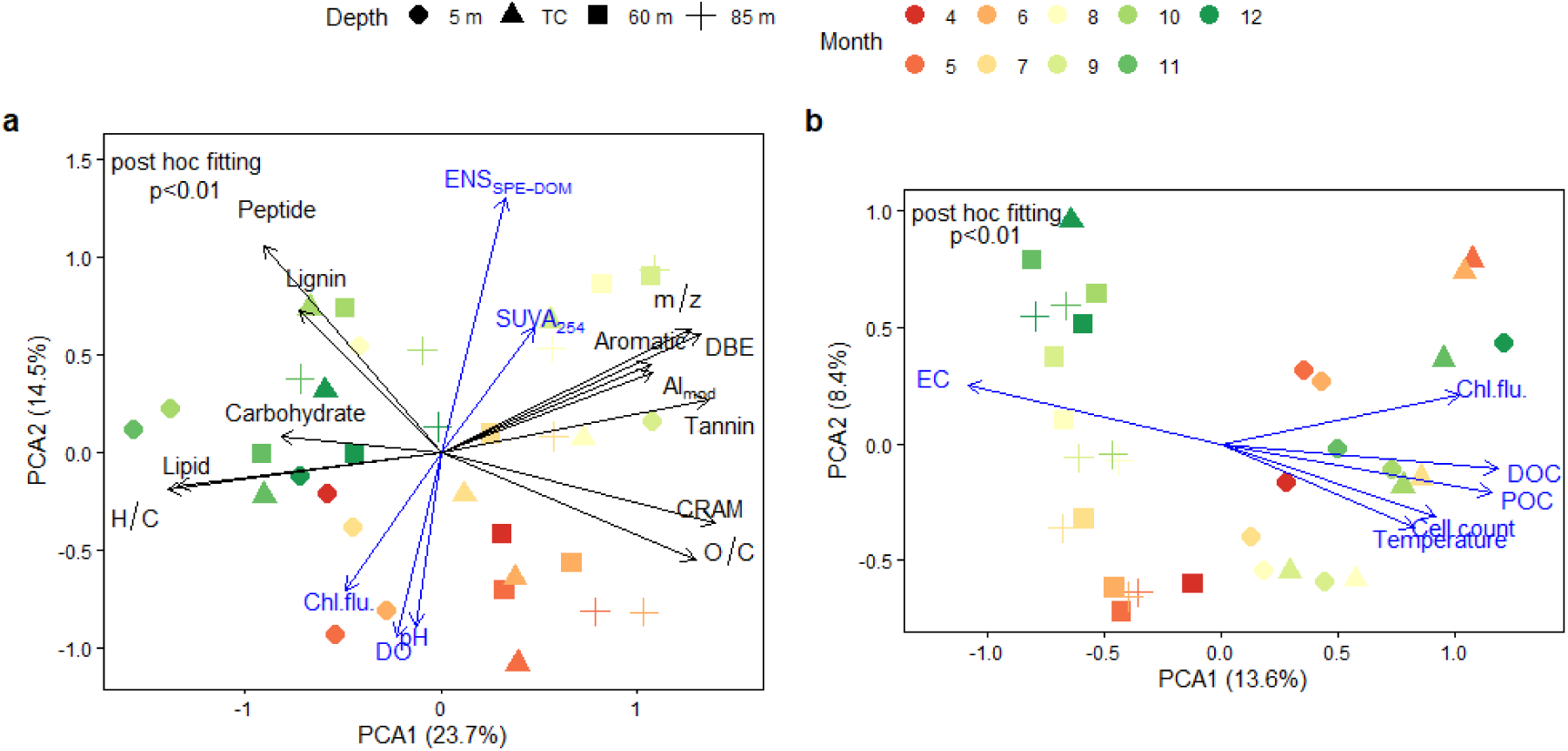
Principal component analysis for SPE-DOM (a) and microbiome (b) relative abundance. Each relative abundance was robust centered logratio-transformed before PCA. External variables were fitted *post-hoc* on the 2D ordination space, where black color indicates average molecular properties and blue color indicates environmental parameters. m/z = mass-to-charge ratio, AI_mod_ = modified aromaticity index, DBE = double-bond equivalent, CRAM = carboxyl-rich alicyclic molecules, ENS_SPE-DOM_ = effective number of species of solid phase-extracted dissolved organic matter, SUVA_254_ = DOC-specific UV absorbance at 254 nm, DO = dissolved oxygen, EC = electrical conductivity, Chl.flu. = chlorophyll fluorescence.

### 2.2. Microbiome

Using 16s rRNA amplicon sequencing, 1541 unique ASVs were detected (113–581 per sample, after removing singlets), with 5 ASVs being common to all samples. The phylogenetic class (and order) of the common ASVs included *Actinobacteria* (*Frankiales*), *Alphaproteobacteria* (*SAR11 clade*), *Kapabacteria* (*Kapabacteriales*), *Verrucomicrobiae* (*Pedosphaerales*), and *Gammaproteobacteria* (*Burkholderiales*). The rank abundance curve of the microbiome was similar to that of SPE-DOM, indicating dominance by a small percentage of abundant ASVs (Fig. 2a–d). Species accumulation curves for microbiome indicated that the detected number of ASVs from different depths was in the order thermocline (TC) > 60 m ≈ 85 m > 5 m (Fig. 2f). The lowest number of species at 5 m was contrary to the highest number of cell abundance at that depth (Fig. 1c), suggesting dominance and proliferation of smaller number of species at the shallowest layer. Microbial diversity quantified as ENS varied by a factor of almost three (18–52), but fluctuated and did not exhibit perceivable seasonal patterns, unlike SPE-DOM (Fig. 3). Within our analytical window, the diversity of SPE-DOM evaluated as MF (i.e., without considering isomers) was one-order larger compared to that of microbiome, highlighting vast and unexplored chemical complexity of organic matter.

Vertical partitioning of the bacterial communities in the epi- and hypolimnion was observed (Fig. S2). At the class-level phylogenetic resolution, *Cyanobacteriia* was relatively more abundant in the epilimnion, while *Anaerolineae*, *Planctomycetes*, *OM190*, and *Nitrospiria* were more dominant in the hypolimnion. There were also some abundant ASVs irrespective of sampling depth, such as *Actinobacteria*, *Bacteroidia*, *Alpha*/*Gammaproteobacteria*, and *Acidimicrobiia* (Fig. S2).

Contrary to SPE-DOM, microbial composition was first separated by depth and secondarily by season (Fig. 4b). These sample distributions were correlated with most environmental parameters and reflected the difference in water chemistry between the epi- and hypolimnion. The first principal component axis separated epilimnetic samples from hypolimnetic samples, while hypolimnetic samples distributed along the second principal component axis with clear seasonality. Seasonality of epilimnetic samples was less clear).

### 2.3. Association between SPE-DOM and microbiome

Canonical correlation analysis (CCorA) found a significant association between the overall compositions of SPE-DOM and microbiome in both the epi- and hypolimnion (p < 0.0001). The first two canonical axes exhibited strong correlations, with values of 0.998 and 0.991 for axis 1 (CCorA1) and 0.987 and 0.932 for axis 2 (CCorA2) in the epi- and hypolimnion datasets, respectively (Fig. S3). However, bimultivariate redundancy coefficients indicated that the explanatory power of SPE-DOM on the microbiome was low in the epilimnion dataset (0.28) (Fig. S3a). In contrast, in the hypolimnion dataset, the explanatory powers of SPE-DOM on microbiome (0.63) and microbiome on SPE-DOM (0.45) were substantially higher (Fig. S3b). Strong canonical correlations do not necessarily mean that the corresponding vectors of ordination scores (canonical variates) explain a large fraction of the variance in both data sets (32). Given this limitation and the original research focus, subsequent analyses were centered on the hypolimnion dataset. In the CCorA 2D multivariate space for the hypolimnion, both SPE-DOM and the microbiome exhibited a clear seasonal pattern (Fig. S3b). Specifically, along CCorA1, sample distribution followed a temporal gradient from the onset of water stratification (April) to just before its breakdown (November and December), transitioning from the positive to negative end of CCorA1 (Fig. S3b).

The covarying hypolimnetic MFs and ASVs were first identified by their common correlations with the first canonical axis (CCorA1). The MFs and ASVs of the same color in Fig. S4 were considered covaried in the hypolimnion of Lake Biwa during stratification due to their common correlations with CCorA1. The MFs significantly correlated with CCorA1 (n = 150) were represented in the Van Krevelen diagram (Fig. S4a). Positively correlated MFs were mostly CHO MFs with O/C ≥ 0.5, while the negatively correlated were mostly CHO or CHON MFs with O/C < 0.5 (Fig. S4a). Significantly correlated ASVs (n = 11) were represented in the phylogenetic cluster (Fig. S4b). Among these, four ASVs were positively correlated and seven were negatively correlated, all of which were affiliated as bacteria (Fig. S4b). At the class level, the significantly correlated bacterial taxa included *Bdellovibrionia*, *bacteriap25*, *Planctomycetes*, *Gammaproteobacteria*, and *Verrucomicrobiae* (Table S3).

The network analysis based on the proportionality index of parts (PIP) between ASVs and MFs identified a much larger number of ASV-MF pairs than CCorA (Fig. 5). In the epilimnion samples (Fig. 5a), three clusters were evident with 639 MFs connected with ASV_00005 and ASV_00026, both belonging to the family *Burkholderiales* within the class *Gammaproteobacteria*. In contrast, the networks in the hypolimnion samples were considerably more complex, involving 19 ASVs and 1705 MFs out of 1755 common MFs, representing the majority of total peak intensities (Fig. 5b). Although small in number, these ASVs accounted for 23 ± 8% of the total relative abundance in the hypolimnion dataset. Notably, the ASVs identified through network analysis differed from those detected by CCorA (Fig. S4). ASV_00162 was affiliated with chloroplast, the organelles responsible for photosynthesis in eukaryotic phytoplankton. Chloroplast sequences are typically considered contaminants in 16S rRNA gene amplicon sequencing and are not of primary interest in quantitative analyses. Furthermore, chloroplast ASVs in the hypolimnion presumably reflect sinking phytoplankton-derived particles, as photosynthesis is not expected in the deep aphotic zone. Nevertheless, with these limitations in mind, we retained chloroplast-associated ASVs in our analysis, as they provide insights into MFs excreted by, or leached from sinking particles of, phytoplankton in a broad context of network analysis. Due to the ratio-based approach and subcompositional coherence of PIP (33), the presence or absence of chloroplast ASVs did not influence the relative abundance inferences of other ASVs, ensuring that network structures among non-chloroplast ASVs and MFs remained unaffected. ASV_00162 exhibited covariance with the highest number of MFs across all compound categories, highlighting the chemical diversity of phytoplankton-derived compounds (Fig. 5b).

**Fig. 5.**
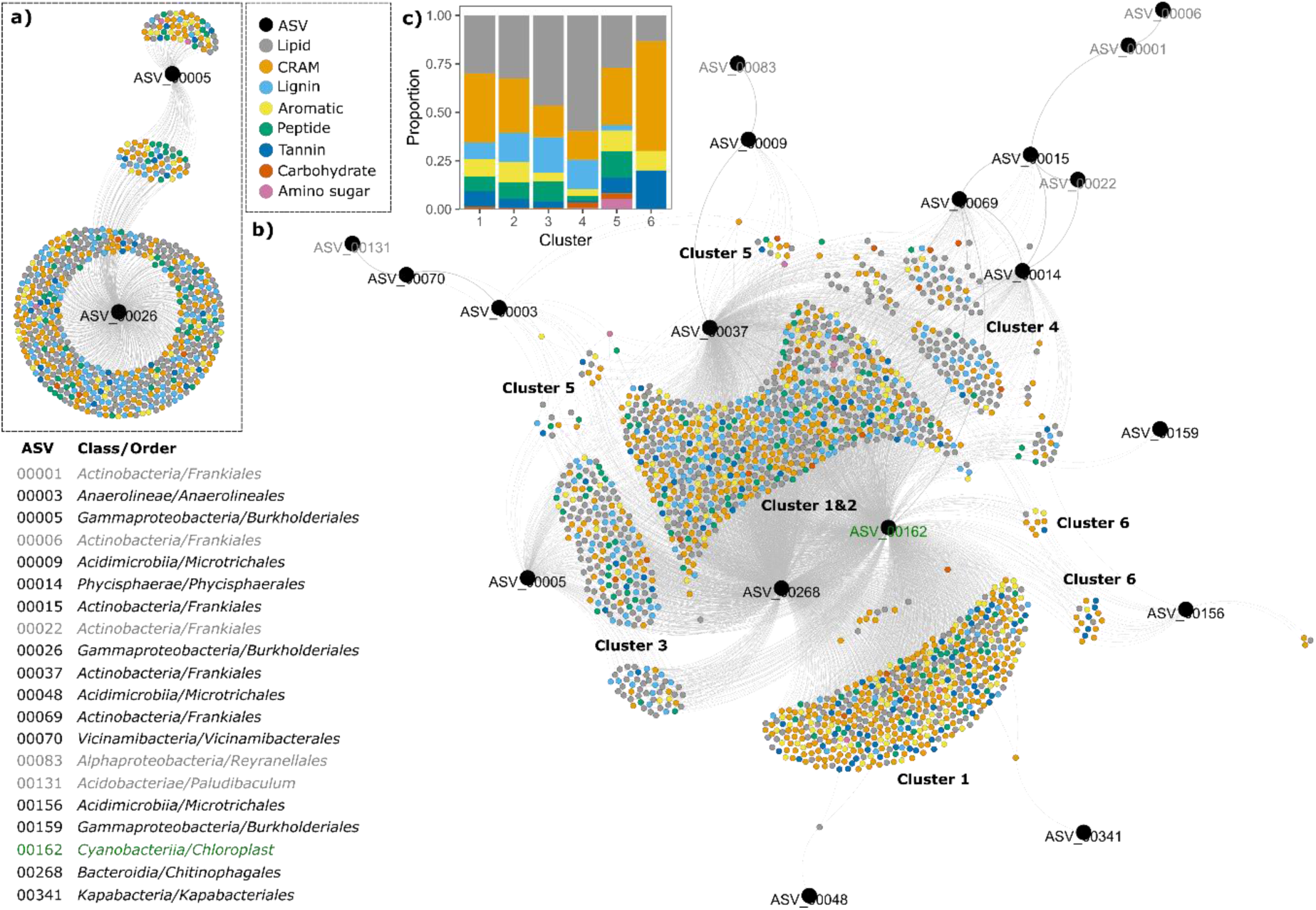
Network of molecular formulae and ASVs in (a) the epilimnion and (b) the hypolimnion during lake water stratification. The positive network was constructed by the proportionality index of parts (PIP). Molecular formulae are color-coded by eight compound categories assigned by the integrated compound category classification (IC3), while ASVs are shown in larger black circles. Six clusters were identified by the modularity class analysis of the networks, and cluster numbers are appended next to each cluster (note, some clusters are separated into several “islands”; see Fig. S5 for the networks color-coded by cluster). The proportion of the compound category in each cluster is shown in (c) in descending order with the same color coding as the network. The class and order of each ASV are presented after the corresponding ASV numbers. Grey ASVs are those without direct linkage with any molecular formula, and a green ASV is a chloroplast (see main manuscript for discussion).

The modularity analysis of the hypolimnion networks detected six clusters (Figs. 5b, S5). The MFs assigned to different clusters tended to be assigned to distinct compound categories (Fig. 5c), which were mirrored in their molecular properties (Fig. S6). Especially, MFs in the smallest clusters (5 and 6) exhibited distinct properties, such as higher O/C ratios and nominal oxidation state of carbon (NOSC) (Fig. S6b, d). Additionally, cluster 6 was associated with MFs having higher m/z (i.e., molecular weight) and DBE, along with lower H/C ratios than other clusters (Fig. S6a, c, e). MFs assigned in the largest clusters (1 and 2) had similar average molecular properties (Fig. S6). However, the proportions of compound categories differed, with more CRAM and fewer Lignin MFs in cluster 1 than cluster 2 (Fig. 5c). CRAMs, enriched in carboxyl functional groups, were highly oxygenated, resulting in relatively higher NOSC values in clusters 1, 5, and 6 than MFs in other clusters (Fig. S7). Conversely, clusters 3 and 4, which were enriched in lipids (Fig. 5c), were characterized by MFs with low NOSC values (Fig. S7).

## 3. Discussion

### 3.1. Seasonality and depth-partitioning of dissolved organic and microbial communities

DOC concentrations and microbial abundance showed stronger seasonality in the epilimnion than in the hypolimnion (Fig. 1). This seasonality in DOC aligns with findings from multiple monitoring studies conducted in the North basin of Lake Biwa between 2001 and 2014 (21, 28, 29). The microbial cell count data are also in good agreement with prior research in this lake (22). Temporal decoupling between DOC peaks and microbial abundance in the epilimnion suggests a lag between photosynthetic DOC production and heterotrophic microbial proliferation. Both spatial and temporal decoupling of DOC production and remineralization have been documented in this lake through vertical and seasonal variations in DOC concentrations and stable isotope signatures (21, 29). Maki et al. (2009) observed the accumulation of ^13^C-rich, semi-labile autochthonous DOC in the epilimnion (estimated δ^13^C: −22.2 ± 0.3‰), which was transported to the hypolimnion during winter overturn and selectively decomposed during stratification. Although this previous study focused on bulk DOM, it provides a broad picture of DOM cycling in this lake, offering insights into the timing, location, and mechanisms of DOM production and degradation relevant to our findings.

PCA indicated that SPE-DOM composition was clearly influenced by seasonality, while microbial composition differed by sampling depth as well as seasonality (Fig. 4). Previous studies have reported clear seasonal variations in fulvic acids in this lake, with more aliphatic signatures in spring and autumn and higher aromaticity (lower H/C) in summer (34), corroborating our findings (Fig. 4a). In contrast, our study found minimal depth-wise variation in SPE-DOM molecular composition, which differs from earlier reports indicating the accumulation of protein-like fluorescence in the epilimnion during stratification (30). This inconsistency may be attributed to differences in analytical windows; our FT-ICR MS analysis of SPE-DOM may have missed large portions of small highly hydrophilic compounds (e.g., hydrophilic amino acids, sugars, and small organic acids) and high molecular-weight compounds (e.g., polysaccharides and proteins) due to the use of PPL (S1). Nevertheless, reported vertical and seasonal gradients in DOC concentration and δ^13^C-DOC suggest the dominacen of a refractory DOM (RDOM) fraction in this lake, which persists on timescales exceeding one year, escaping decomposition in the epilimnion during stratification. This RDOM fraction accounted for >90% of bulk DOM during the overturn period (29), explaining the minimal depth variation observed in our analysis (Table 1, Fig. 4a), which effectively captured this refractory component.

The vertical partitioning of bacterial communities in this lake has been well documented (22). At the class level, we found that *Anaerolineae*, *Planctomycetes*, *OM190*, and *Nitrospiria* were predominant in the hypolimnion (Fig. S2). These bacteria were all identified as the hypolimnion habitat specialists (22). About seasonality?? Given the scarcity of research regarding the bacterioplankton inhabiting the hypolimnion of deep freshwater holomictic lakes, our interests in this study were the association between these hypolimnetic specialists and specific MFs which can give valuable insights into their ecology and ecophysiology.

### 3.2. Tight coupling between dissolved organic matter and bacteria in the oxygenated hypolimnion of a deep freshwater lake

The network analysis uncovered far more than simple associations: it disentangled a molecular– microbial web of interactions that underpins the transformation and persistence of DOM in deep waters.(Fig. 5). This analysis, which was based on ASV-MF level associations, independently evaluated the proportionality of each ASV or MF (33). Consequently, the network was not influenced by the overall multivariate patterns, as suggested in CCorA (Supplementary Discussion S3), and the network analysis captured more nuanced relationships between ASVs and MFs. Furthermore, this network analysis employed a robust and valid association measure for compositional data between ASVs and MFs, providing coherent results independent of the considered subsets of data or normalization methods (17, 33, 35). The stark contrast in network complexity between the epi- and hypolimnion corresponds with the CCorA results, where MFs accounted for a substantial portion of ASV variation only in the hypolimnion (Fig. S3a), highlighting the importance of the coupling between DOM and microbes in the deep (discussions on the epilimnion network can be found in S3). Furthermore, water stratification of Lake Biwa throughout the study period minimized influences from co-factors such as water mass mixing or terrestrial inputs on the hypolimnion which can produce apparent correlations between ASVs and MFs (9). Collectively, the identified networks likely represent real ASV-MF associations that were not primarily influenced by a common environmental factor.

The hypolimnion network analysis revealed that hypolimnion bacteria were associated with MFs in different clusters (Fig. 5b). Among the ASVs in the hypolimnion network (Fig. 5b), ASV_00003 (CL500-11, phylum *Chloroflexi*), ASV_00014 (CL500-3, phylum *Planctomycetota*), and ASV_00341 (a member of Kapabacteria) were hypolimnion specialists (22). So far, the substrate preferences and potential symbiotic relationships of these bacteria remain unidentified. ASV_00003 was associated with cluster 5 MFs, ASV_00014 with cluster 4 MFs, and ASV_00341 with cluster 1 MFs (Fig. 5b). All these clusters contained diverse MFs covering almost all compound categories, though the relative abundance of compound categories differed among clusters. Cluster 5 was enriched in labile compounds such as Amino sugar, Carbohydrate, and Peptide MFs (Fig. 5c). Cluster 4 was dominated by Lipid MFs (Fig. 5c). Cluster 1 had the largest contribution from CRAM MFs but also contained relatively high proportions of Tannin MFs, both of which are considered relatively refractory (Fig. 5c). These distinct substrate preferences of hypolimnetic bacteria suggest their broad ecological niche. Of particular interest is CL500-11 (phylum *Chloroflexi*), which has been recently reported as globally ubiquitous in oxygenated hypolimnia of deep freshwater lakes, where it likely plays a crucial role in material cycling (23, 25, 26). Based on available metagenomic and geochemical data, CL500-11 is thought to preferentially utilize autochthonous, microbial-derived N-rich DOM produced in the hypolimnion, which presumably turnovers within an annual timescale, such as di-and oligopeptides, spermidine, putrescine, branched-chain amino acids, and N-acetylglucosamine (25, 26). Our network analysis supports this proposed substrate preference, with CL500-11 showing associations with labile, N-rich Amino sugar and Peptide MFs, providing another line of evidence from the environmental metabolomics standing point.

The remarkable network complexity suggests that hypolimnion specialists not only exploit diverse classes of DOM molecules but also engage in complementary roles. Among the hypolimnion specialists, CL500-11 and ASV_00014 formed links with other ASVs (Fig. 5b). CL500-11 co-occurred with ASV_00070 (class *Vicinamibacteria*) which, in turn, co-occurred with ASV_00131 (*Acidobacteriae*), both within the phylum *Acidobacteriota* (Fig. 5b). *Acidobacteria* are known for their ability to decompose a range of complex substrates, including hemicellulose, cellulose, chitin, and aromatic compounds, to produce smaller compounds (36, 37) Therefore, CL500-11 may benefit from the decomposition byproducts produced by *Acidobacteriota*. ASV_00014 co-occurred with three ASVs 00069, 00022, and 00015 which also co-occurred with ASV_00001 and ASV1_00006 (Fig. 5b). ASV_00014 is a member of the class *Phycisphaerae* of the phylum *Planctomycetota* and is considered to be capable of utilizing phytoplankton-derived high molecular weight organic matter rich in polysaccharides (38). Notably, all but one of these ASVs were affiliated with the identical *hgcI clade* of phylum *Actinobacteriota* (Table S4). The *hgcI clade (acI clade)* has been reported as a dominant fraction of bacterioplankton in freshwater lakes (38, 39) and has a strong genetic ability to use carbohydrate and N-rich organic compounds (38, 40). Most of these ASVs did not show links with MFs in cluster 4, which was enriched in more recalcitrant Lipid MFs rather than carbohydrate and N-rich organic compounds (Fig. 5b). This suggests that ASV_00014 may function as a hub, providing decomposition byproducts from Lipid MFs breakdown to these ASVs. The co-occurrence of these taxa and their shared compound clusters point to potential metabolic cooperation and symbiotic exchange of decomposition products, revealing how microbial communities collectively sustain carbon cycling in the deep.

Finally, these substrate preferences of hypolimnetic bacteria associated with clusters 4 & 5 may be explained by the energetics associated with the oxidative degradation of organic matter. MFs in cluster 5 had higher NOSC values than those in cluster 4 (Fig. S6d), corresponding with lower Gibbs energies for the oxidation half reactions of organic compounds (41). On average, the removal of an electron from an organic compound (i.e., oxidation) becomes thermodynamically more favorable as NOSC increases, thus such compounds tend to be microbially labile (41). CL500-11 (associated with cluster 5) likely prefers compounds that are thermodynamically more readily used. Contrarily, ASV_00014 (associated with cluster 4) is likely capable of utilizing organic compounds with lower NOSC values that are thermodynamically more difficult to oxidize (Fig. S6d, S7). The existence of such hub bacteria with a high oxidation capability may play a substantial role in the proliferation of major bacteria such as the *hgcI clade*.

Overall, this study provides the first comprehensive investigation of the interactions between DOM and microbial communities in the epilimnion and hypolimnion of a deep freshwater lake. We demonstrated that the hypolimnion harbors more complex DOM-microbe networks, highlighting stronger coupling between microbial communities and organic substrates during stratification. The networks can be viewed as molecular–microbial blueprints of organic matter decomposition and production, capturing the ecological strategies by which microorganisms regulate the fate of DOM under stable stratified conditions. Such molecular–microbial blueprints have implications that extend well beyond Lake Biwa. They provide a conceptual and methodological leap forward in linking microbial ecology with environmental metabolomics, setting the stage for predictive models of organic carbon cycling across diverse aquatic ecosystems, from freshwater lakes to the ocean interior. Future studies could focus on several key areas to build upon the findings of this research, including longitudinal studies for longer years enabling causal inference and characterization of microbial functional roles in DOM decomposition and transformation involving metagenomic and metatranscriptomic approaches to examine microbial genes and metabolic pathways involved in the degradation of various DOM fractions. Integrating biogeochemical modeling with metabolomic and microbiome data would enable predictions of how changes in DOM composition and microbial communities may impact broader ecosystem functions such as carbon and nutrient cycling.

## 4. Materials and Methods

### 4.1. Sampling

We collected water samples monthly from the epilimnion (5 m, thermocline (TC)) and hypolimnion (60 m, 85 m) at a pelagic station (station 17B: 35°23.5’ N, 136°07.3’ E; water depth: ca. 89 m) of the north basin of Lake Biwa, Japan, from April to December 2022 (Table S1). In April 2022, we collected water samples only at 5 m and 60 m. Lake Biwa has a surface area of 670 km^2^, a water storage capacity of 275 km^3^, a maximum depth of 104 m, and an estimated water residence time of approximately 5.5 years. Lake Biwa consists of two basins, a small shallow South basin and a large deep North basin. The main North basin is mesotrophic and monomictic and clearly stratified from April/May to December, when the hypolimnion remains oxygenated. Therefore, our sampling aimed to investigate DOM-microbe dynamics from the onset to the break of lake water stratification.

We measured the vertical profiles of basic water properties (Temperature, electrical conductivity, chlorophyll fluorescence, turbidity, and dissolved oxygen (DO)) *in situ* using a CTD profiler (AAQ-RINKO AAQ176, JFE Advantech, Japan) (Table S1). We collected water samples directly from an autonomous water sampler with 5-L Niskin bottles (AWS387-5, JFE Advantech, Japan). We stored the samples in 1 L, detergent and acid-washed polypropylene bottles after rinsing them twice with the sampled water, which was then stored in a cool box immediately after sampling and transported to the laboratory. To remove large particles such as zooplankton during sample collection, we used a 250 μm mesh stainless steel sieve as prefiltration during water collection. We filtered samples within eight hours after sample collection using a 0.1 μm pore size hydrophilic PTFE membrane filter (47 mm diameter: Omnipore JVWP04700, Merck). We immediately froze aliquots of the filtrates and filters and stored them at –28°C until optical analysis and DNA extraction, respectively. For sample preservation for solid phase extraction, we immediately acidified filtrates with 1M HCl to pH 2 and stored them in the fluorinated polypropylene bottles in the dark at 4°C. For cell counting, we fixed the prefiltered water (< 250 μm) with 1% (final concentration) glutaraldehyde (20% for electron microscopy; Fujifilm Wako Pure Chemical) within five hours after sample collection and stored them at 4°C for one hour. We then quickly froze the samples for cell counting in liquid nitrogen and stored them at −80°C.

### 4.2. Chemical analysis

We measured the concentrations of total organic carbon (TOC) and dissolved organic carbon (DOC) using a TOC analyzer (TOC-L_CPN_ for monitoring a microscopic algae biomass, Shimadzu, Japan) within the day of sampling. For TOC analysis, we treated samples in an ultrasonic bath for 10 min and immediately measured. We measured DOC in three different vials and TOC in three vials for surface water (5 m and TC) and six vials for deep water samples (60 m and 85 m). Particulate organic carbon (POC) was estimated as the difference between TOC and DOC. The analytical precision (standard error) among vials was on average 0.01 mgC L^-1^ for DOC and 0.02 mgC L^-1^ for TOC and POC, respectively (Table S1). We collected UV−vis absorbance spectra (239−560 nm) using a UV−visible spectrophotometer (Aqualog-UV-C, Horiba, Japan) with a 1 cm path length quartz cuvette. Using DOC and UV-vis data, we calculated the spectral slope (S_275−295_, in nm^−1^) and DOC-specific UV absorbance at 254 nm (SUVA_254_, in L mg C^−1^ m^−1^) as previously reported (42, 43) (Table S1).

### 4.3. FT-ICR MS analysis

We extracted and desalted DOM for FT-ICR MS using Agilent Bond Elut PPL (100 mg) cartridges following a protocol described earlier (44) with a slight modification regarding cartridge preparation and DOC loading (details in Supplementary Method S1). The average extraction efficiency was, on average, 51% ± 5% on a DOC basis (Table S1). The extracted DOM, hereafter referred to as solid-phase extracted DOM (SPE-DOM), was within the analytical window of FT-ICR MS. We performed mass spectrometric analysis of SPE-DOM on a 7 Tesla solariX FT-ICR mass spectrometer (Bruker Daltonik, Bremen, Germany), equipped with an electrospray ionization source (ESI, Bruker Apollo II) applied in the negative ionization mode (details in S1). The instrumental stability and inter-laboratory comparabilty were verified using a reference material purchased from the International Humic Substances Society (Suwanee River Natural Organic Matter, SRNOM; 2R101N), yielding results consistent with reported reference values (45) (Table S2). We assigned MFs of detected masses as previously described (44, 46) using an online mass spectrometry data analysis pipeline (47). In total, we used 6886 MFs for further analysis after deleting singlets (i.e., MFs detected in only one sample), isotopologues, and those with double bond equivalents (DBE) minus the number of O atoms of greater than 10 (48). No peaks occurred above 600 m/z. Besides DBE, molecular indices such as modified aromaticity index (AI_mod_) (49) and nominal oxidation state of carbon (NOSC) (41) were calculated.

We grouped MFs into nine compound categories based on the combination of best available classification rules, including multidimensional stoichiometric compound classification (MSCC) of biomolecules using C:H:O:N:P stoichiometries (50), NMR data-based assignments of lignin and tannin (51), DBE-to-elemental ratios for carboxyl-rich alicyclic molecules (CRAM) (52), and the unequivocal criterion of aromatic structures by AI_mod_ (49): lipids, peptides, amino sugars, carbohydrates, nucleotides, lignin, tannin, aromatic, and CRAM. CRAM are considered a major refractory component of natural organic matter and oxidative decomposition products of biomolecules (52). The superiority of Rivas-Ulabch’s “beyond the van Krevelen Diagram” MSCC approach over existing methods was thoroughly explained in their papers, but MSCC is developed for biomolecules and is not necessarily suitable for environmental research (Rivas-Ubach et al., 2018). By further tailoring for natural organic matter incorporating lignin, tannin, and CRAM, our integrated compound category classification (IC3) is arguably the most comprehensive and unambiguous classification rule for environmental research (details in Supplementary Method S2). By using this rule, we were able to assign compound categories to 6864 MFs out of 6886 MFs (no nucleotides assigned), achieving 99.7% assignment (Fig. S1). Similarly, we categorized MFs based on the containing elements into compound classes, namely, CHO, CHON, CHOS, CHOP, CHONS, CHONP, and CHOSP. Finally, we calculated the relative abundance of each category and average molecular properties (H/C, O/C, m/z, AI_mod_, and DBE) per sample as weighted averages considering FT-ICR MS relative peak intensities. Because compound categories based on the elemental stoichiometry are not proof of the existence of such compounds, we distinguish them by the capitalized terms, such as Amino sugar, Aromatic, and so forth.

### 4.4. Microbial analysis

For cell counting, we stained cells in the glutaraldehyde-fixed samples with 1× SYBR Green I (Thermo Fisher Scientific) and analyzed them using a flow cytometer (Beckman Coulter, CytoFLEX) equipped with a 488 nm excitation laser. For each sample, we measured the side scatter, green (525 nm), and orange (690 nm) fluorescence and determined the number of stained cells on the cytogram. Based on the flow rate of the analysis, we calculated the bacterial cell density in the original water sample.

For 16S rRNA gene amplicon sequencing, we extracted DNA from the filters using DNeasy PowerSoil Pro Kit (Qiagen) following the manufacturer’s protocol (details in Supplementary Method S1). The extracted DNA was quantified using Qubit 4 fluorometer (Thermo Fisher Scientific) and amplified using universal primer sequences for prokaryotes, 515F-Y (5′-GTGYCAGCMGCCGCGGTAA) and 926R (5′-CCGYCAATTYMTTTRAGTTT) (53). The resulting sequencing library was input to Illumina NovaSeq 6000 pair-end (2 × 250 bp) sequencing, and we generated at least 16,000 (average: 68,000) sequence pairs per sample. We used DADA2 v1.30.0 (54) in R v4.3.2 (http://www.R-project.org/) to process the demultiplexed raw sequencing reads following the general workflow of the software (details in Supplementary Method S1). We assigned taxonomy to each ASV with the DADA2-formatted SILVA NR99 version 138 database (55). ASVs detected in only one sample (singlets) were removed. Finally, we combined the ASV table with the taxonomic assignment and transformed the read count to relative abundance by dividing the counts by the total read number of each sample.

### 4.5. Statistical analysis

#### 4.5.1. Community structure and ordination analysis

We conducted all statistical analyses using R. The rank abundance curves and species accumulation curves for SPE-DOM and 16s rRNA-based microbiome were drawn. As indicators for alpha diversity (i.e., within-sample diversity), we calculated the effective number of species (ENS) (56) for both SPE-DOM and microbial communities, converted from the Gini-Simpson index based on the relative peak intensities of FT-ICR MS spectra. ENS contains both richness and abundance information and has a “doubling” property, making it an intuitive measure of diversity that is linearly comparable among communities (56).

We performed principal component analysis (PCA) of the FT-ICR MS relative peak intensities and the 16S rRNA gene-based ASV relative abundance to summarize their major variation. Given the *compositional* nature (17) of both FT-ICR MS relative peak intensities and ASV relative abundance, a robust centered logratio (rclr) transformation (57) was applied before PCA. External environmental variables and intensity-weighted average molecular properties were fitted *post-hoc* to the PCA ordination spaces with p-values calculated over 9999 permutations.

#### 4.5.2. DOM-microbe association

We used canonical correlation analysis (CCorA) to find community-level covariation between chemical (SPE-DOM) and microbial assemblages. CCorA is a multivariate symmetrical analysis that finds linear combinations of variables that maximize correlation between two corresponding data matrices. We performed two separate CCorAs for the epi-(5m and TC) and hypolimnion (60 m and 86 m) datasets, given the distinct microbial composition in these two water layers (Fig. 4b). Details of CCorA are provided in Supplementary Method S1.

We then performed network analysis at the ASV-MF resolution using the proportionality index of parts (PIP) (33). Unlike ordinary correlation metrics, which are unsuitable for compositional data due to their dependence on data subsetting or normalization (17, 35), PIP explicitly utilizes ratios between components, enabling robust inference of associations within and between MFs and ASVs. Since ratios remain invariant to subsetting or normalization, ratio transformation serves as a fundamental step in compositional data analysis. Among several compositional metrics, only PIP remains constant irrespective of the considered set of variables (i.e., subcompositionally coherent), making it the most robust index to study ASV-MF association (33). To construct the core network, we used common MFs and ASVs detected in more than half of the samples and analyzed the network separately for the epi- and hypolimnion. As ratio calculations are not possible in the presence of zeros, we substituted zeros in the ASV data with 1/10^th^ or 1/100^th^ of the smallest nonzero value in the dataset. We retained MF-ASV pairs and ASV-ASV pairs with PIP > 0.75 and visualized the resulting networks using the open-source software Gephi (https://gephi.org/). Zero substitution at either threshold yielded consistent results, preserving 691 ASV-MF links for the epilimnion and 5167 ASV-MF or ASV-ASV links in the hypolimnion. Isolated ASV-ASV links without connections to any MF were subsequently removed, and retained MFs were color-coded by either compound categories or NOSC. Finally, we identified clusters using modularity class analysis and calculated average molecular properties of MFs within each cluster.

## Data availability

The raw sequencing reads generated in the present study were deposited under the BioProject accession number PRJDB20674. FT-ICR MS data, ASV table and sequences, environmental data, and network data (nodes and edges files for Gephi) are freely available at figshare: 10.6084/m9.figshare.28667366.

## Author contributions

MK conceived the study. YTY, ASG, and KH conducted sample collection and processing. AO and MK analyzed DOM samples by FT-ICR MS with significant support from HN, YO analyzed 16S rRNA amplicon sequence data, KH conducted the *in-situ* water profiling, and YTY analyzed other basic water chemical parameters. AO and MK conducted all statistical analyses. MK wrote the initial draft. All authors participated in finalizing the manuscript.

CRediT roles:

Conceptualization, MK; Formal analysis, AO, MK; Funding acquisition, MK, YO, YTY, HN; Investigation, AO, MK, YO, YTY, ASG, KH; Methodology, MK, YO, YTY, HN; Project administration, MK; Resources, MK, YO, YTY, KH, HN; Supervision, MK; Visualization, AO, MK; Writing – original draft, MK; Writing – review & editing, YO, YTY, ASG, KH, HN

## Competing Interests

The authors declare no competing interests.

## Acknowledgements

We thank the captain and crews of R/V Biwakaze for helping with the sample collection. Additional thanks to M. Morikawa, T. Uenishi, and H. Amimoto in LBERI, and TT. Nguyen and S. Okuda in Kyoto University for their assistance with sample collection, preparation, and chemical analysis. This work was supported by JSPS KAKENHI [grant numbers JP22H03733/JP23K24987 (MK), JP22K15182 (YO), JP21H03584 (YTY), JP22H03723/JP23K24977 (MK, YTY, KH), and JP22H00382 (YO, YTY, KH)] and JST FOREST (Fusion Oriented REsearch for disruptive Science and Technology) Program [grant numbers JPMJFR231C (MK) and JPMJFR2273 (YO)]. The access to FT-ICR MS was provided by the Analysis and Development System for Advanced Materials (ADAM) collaboration program of RISH, Kyoto University. MK was also partially supported by startup funding from Kobe University.

## Notes

https://doi.org/10.6084/m9.figshare.28667366.v1

### Competing Interest Statement

The authors have declared no competing interest.

