## Supporting Information for "Couples in the deep: dissolved organic and microbial communities in the oxygenated hypolimnion of a deep freshwater lake"

**Contents:**

Supplementary Method S1

Supplementary Method S2

4 Supplementary Tables

7 Supplementary Figures

**Supplementary Method S1.**

S1-1. SPE-DOM extraction

We extracted and desalted DOM for FT-ICR MS using Agilent Bond Elut PPL (100 mg) cartridges (Dittmar et al., 2008) with a slight modification regarding cartridge preparation and DOC loading. We filled the cartridges with methanol (Guaranteed Reagent, FUJIFILM Wako Chemicals, Japan), kept them overnight, and drained them. Then, we blew out residual methanol in the resin by pushing air with a plastic syringe and rinsed it three times with pH 2 ultrapure water. To prevent overloading, we loaded the sample volume corresponding to 0.24 mgC (Li et al., 2016). After loading samples, we rinsed the cartridges three times with pH 2 ultrapure water, dried with $N_2$ gas, and eluted with methanol. We stored the methanol extracts in 2mL, acid-washed and muffled amber vials at −20 °C in the dark until analysis. The average extraction efficiency was, on average, 51% ± 5% on a DOC basis (Table S2). To check for possible contaminations, we prepared an operational blank extract by processing ~200 mL of pH 2 ultrapure water (0.01 M HCl) the same way as the samples at each day of extraction. The extracted DOM, hereafter referred to as solid-phase extracted DOM (SPE-DOM), was within the analytical window of FT-ICR MS.

The analytical window in this study should be further elaborated to clarify the potential and limitations of our approach in studying DOM-microbial associations; the PPL cartridge used to extract SPE-DOM is a styrene-divinylbenzene polymer modified with a nonpolar surface, with a mean diameter of 125 µm and pore size of 150Å (15 nm). Due to its hydrophobic property, PPL preferentially extracts hydrophobic compounds from the bulk DOM. However, its selectivity for hydrophobic compounds is less clear compared to that of classical hydrophobic resins such as XAD-8 (or DAX-8) and C18 (Li et al., 2017; Perminova et al., 2014). The pore size and tightly packed sorbents of PPL likely retain high-molecular-weight colloidal compounds at the top of the cartridge, preventing their inclusion in SPE-DOM extracts (Hawkes et al., 2016). Chemically excluded compounds (e.g., hydrophilic amino acids, sugars, and small organic acids) and physically excluded compounds (e.g., polysaccharides and proteins) represent some of the most labile DOM fractions, which are readily utilized or decomposed by heterotrophic microbes (Benner and Amon, 2015). Consequently, PPL preferentially retains semi-labile to refractory DOM fractions, which cycle on timescales of >days to years; thus, the DOM-microbe associations in this study target similar temporal scales (>days to the stratification period).

S1-2. FT-ICR MS analysis

We performed mass spectrometric analysis of SPE-DOM on a 7 Tesla solariX FT-ICR mass spectrometer (Bruker Daltonik, Bremen, Germany), equipped with an electrospray ionization source (ESI, Bruker Apollo II) applied in the negative ionization mode. This soft ionization can ionize even high molecular-weight compounds such as proteins without fragmentation but induces a bias that polar compounds with acidic functional groups (such as carboxyls and phenols) are preferentially ionized and detected (Kujawinski et al., 2002; Novotny et al., 2014). We optimized analytical conditions including sample concentration and instrumental parameters regarding ionization and ion transfer, using a reference material purchased from the International Humic Substances Society (Suwanee River Natural Organic Matter, SRNOM; 2R101N). We prepared a SRNOM stock solution by mixing the powder in a methanol:ultrapure water (1:1 v/v) solution, filtering, and storing it in a glass bottle. Before analysis, we diluted the SPE-DOM extracts to a final concentration of 20 mg C $L^{-1}$ in a mixture of methanol and ultrapure water (1:1 v/v). Samples were injected at a rate of 180 $\mu L^{-1}$, the capillary voltage of 3.9 kV, and the drying gas flow rate of 3.7 L $min^{-1}$ at 240 °C. A total of 100 transients with a transient size of 4 mega word (MW) data points in a scanning range of 150–1800 Da were co-added for each sample with a time of flight of 0.65 ms. We optimized ion accumulation time in the hexapole (0.1–0.6 s) for each sample. To test the instrument reproducibility and stability, we analyzed the SRNOM solution of 20 mg C $L^{-1}$ under the identical settings. All samples were analyzed within two days in a random order.

### S1-3. Cell count

We stained cells in the glutaraldehyde-fixed samples with 1× SYBR Green I (Thermo Fisher Scientific) and analyzed them using a flow cytometer (Beckman Coulter, CytoFLEX) equipped with a 488 nm excitation laser. For each sample, we measured the side scatter, green (525 nm), and orange (690 nm) fluorescence and determined the number of stained cells on the cytogram. Based on the flow rate of the analysis, we calculated the bacterial cell density in the original water sample.

### S1-4. 16S rRNA amplicon sequencing

We extracted DNA from the filters using DNeasy PowerSoil Pro Kit (Qiagen) following the manufacturer's protocol. The extracted DNA was quantified using Qubit 4 fluorometer (Thermo Fisher Scientific) and amplified using universal primer sequences for prokaryotes, 515F-Y (5′-GTGYCAGCMGCCGCGGTAA) and 926R (5′-CCGYCAATTYMTTTRAGTTT) (Parada et al., 2016). We performed the early-pooling protocol with two-step PCR to prepare a multiplexed amplicon sequencing library (Ushio et al., 2022). The first PCR to add an index tag to each sample was performed with initial denaturation at 98°C for 30 s, followed by 35 cycles of amplification (denaturation at 98°C for 10 s, annealing at 60°C for 10 s, and extension at 72°C for 15 s) and the final extension at 72°C for 300 s. The PCR products were purified using ExoSAP-IT Express (Thermo Fisher Scientific), quantified, equimolarly pooled, and purified again using AMPure XP beads (Beckman Coulter). The second PCR to add the sequencing adaptors to the pooled amplicons was performed with initial denaturation at 98°C for 30 s, followed by 10 cycles of amplification (denaturation at 98°C for 10 s, and annealing and extension at 72°C for 15 s) and the final extension at 72°C for 300 s. We used Platinum

SuperFi II PCR Master Mixes (Thermo Fisher Scientific) for both first and second PCR. Finally, we purified the second PCR product using AMPure XP beads and E-Gel SizeSelect II system (Thermo Fisher Scientific). The resulting sequencing library was input to Illumina NovaSeq 6000 pair-end (2 × 250 bp) sequencing, and we generated at least 16,000 (average: 68,000) sequence pairs per sample.

#### S1-5. Analysis of sequencing reads

We used DADA2 v1.30.0 (Callahan et al., 2016) implemented in R v4.3.2 (http://www.R-project.org/) to process the demultiplexed raw sequencing reads. Following the general workflow of the software, we subsequentially applied the *filterAndTrim*, *learnErrors*, *dada*, *mergers*, *makeSequenceTable*, and *removeBimeraDenovo* functions to generate an ASV table, a table showing the number of reads assigned to each ASV in each sample. Specifically, we applied "trimLeft=c(19,20), truncLen=c(230,210), maxN=0, maxEE=c(2,2), truncQ=2, rm.phix=TRUE" parameters for *filterAndTrim* function. We assigned taxonomy to each ASV using *assignTaxonomy* function with the DADA2-formatted SILVA NR99 version 138 database (Quast et al., 2012) provided on the software website. ASVs detected in only one sample (singlets) were removed. Finally, we combined the ASV table with the taxonomic assignment and transformed the read count to relative abundance by dividing the counts by the total read number of each sample.

#### S1-6. Canonical correlation analysis

We used canonical correlation analysis (CCorA) to find community-level covariation between chemical (SPE-DOM) and microbial assemblages. CCorA is a multivariate symmetrical analysis that finds linear combinations of variables that maximize correlation between two corresponding data matrices. The symmetrical analysis means that it treats both matrices equally and does not assume a causal relationship (Legendre and Legendre, 2012). Because CCorA cannot handle more variables than observations, we used the first principal coordinates of principal coordinate analysis (PCoA) as inputs to CCorA (Anderson and Willis, 2003), to align with past studies employing the same technique (Osterholz et al., 2016). We performed two separate CCorAs for the epi- (5m and TC) and hypolimnion (60 m and 86 m) datasets, given the distinct microbial composition in these two water layers (Fig. 4b). We used the first principal coordinates of chemical and microbial data explaining more than 75% of the total variance in the Bray-Curtis dissimilarity matrix of each dataset for CCorA (PCoA 1-6 for epilimnion and PCoA 1-5 for hypolimnion). Significance of canonical correlation was computed over 9999 permutations. We calculated bimultivariate redundancy coefficients to evaluate how much of the SPE-DOM composition is explained by the 16S rRNA-based microbiome composition and vice versa. After conducting CCorA, we calculated the Spearman rank correlation coefficient for original variables (i.e., the relative abundance of MFs or ASVs) with the first canonical axis (Anderson and Willis, 2003) to identify the co-varying MFs or ASVs.

For this correlation analysis, we used only MFs and ASVs detected in more than half of the samples. This correlation analysis detects only MFs or ASVs that roughly monotonically varied along the canonical axis. We visualized and color-coded these correlations in a Van Krevelen diagram and a phylogenetic cluster. Only significantly correlated ($p < 0.05$) features were retained after a Benjamini-Hochberg correction for multiple significance testing that can control the false discovery rate.

**Supplementary Method S2.**

Integrated compound category classification (IC3) rule

Worked example in R

```
# Import data -------------------------------------------------------------------
Library(dplyr)

# Molecular formulae (MF) in rows and molecular properties and samples in columns for each MF
# Output from ICBM-OCEAN can be directly used
MF <- read.csv(file = "YourFile.csv", header = TRUE, row.names = 1)

# Remove MF containing minor isotopes; this is optional
# Numbers in comment after command indicate a change in the numbers of MF in the L.Biwa dataset used in this study
MF <- MF[MF$C13 == 0 & MF$N15 == 0 & MF$O18 == 0 & MF$S34 == 0, ] # 9007->7709

# Remove MF with DBE-O >= 10 following Herzsprung et al. (2014); this is optional
MF$DBEminusO <- MF$DBE - (MF$O)
MF <- subset(MF, DBEminusO < 10) # 7709 ->7022

# Remove some unwanted samples (optional)
MF <- MF[-c(113:120)] # remove LB2302-2303

# Remove singlets and MF no longer detected in the remaining samples
MF <- cbind(rowSums(!is.na(MF[79:112])), MF)
MF <- subset(MF, MF[1] > 1)  # 7022 -> 6886
MF[1] <- NULL

# Check number of common MF
MF <- cbind(rowSums(!is.na(MF[79:112])), MF)
dplyr::count(MF, MF[1] == 34)  # 1755
MF[1] <- NULL


# Two main published compound category assignment rules --------------
# Data-based compound category rules by Laszakovits & Mackay (2022).
# Lots of unassigned MF, especially at the center of VK diagram, thus not ideal for NOM research as is.
# amino-sugar (O.C >= 0.56 & O.C <= 0.95 & H.C >= 1.62 & H.C <= 2.35)
# carbohydrate (O.C >= 0.56 & O.C <= 1.23 & H.C >= 1.53 & H.C <= 2.2)
# lignin (O.C >= 0.21 & O.C <= 0.44 & H.C >= 0.86 & H.C <= 1.34)
# lipid (O.C >= 0.01 & O.C <= 0.35 & H.C >= 1.34 & H.C <= 2.18)
# peptide (O.C >= 0.17 & O.C <= 0.48 & H.C >= 1.33 & H.C <= 1.84)
# tannin (O.C >= 0.16 & O.C <= 0.84 & H.C >= 0.7 & H.C <= 1.01)
# But lignin and tannin are useful as these are not considered in Rivas-Ubach et al. (2018).

# The multidimensional stoichiometric constraints classification (MSCC) by Rivas-Ubach et al. (2018).
# The most accurate classification of biomolecules, but not tailored for NOM.
# Much less unassigned MF than Laszakovits & Mackay (2022). Thus, this should be executed first as basis.
# lipid (O.C <= 0.6 & H.C >= 1.32 & N.C <= 0.126 & P.C < 0.35 & N.P <= 5)
# carbohydrate (O.C >= 0.8 & H.C >= 1.65 & H.C < 2.7 & N == 0)
# amino-sugar (O.C >= 0.61 & H.C >= 1.45 & N.C <= 0.2 & N.C > 0.07 & P.C < 0.3 & N.P <= 2 & O >= 3 & N >= 1)
# phytochemical/oxyaromatic compound (O.C <=1.15 & H.C < 1.32 & N.C < 0.126 & P.C <= 0.2 & N.P <= 3)
# protein 1 (O.C > 0.12 & O.C <= 0.6 & H.C > 0.9 & H.C < 2.5 & N.C >= 0.126 & N.C <= 0.7 & P.C < 0.17 & N >= 1)
# protein 2 (O.C > 0.6 & O.C <= 1 & H.C > 1.2 & H.C < 2.5 & N.C > 0.2 & N.C <= 0.7 & P.C < 0.17 & N >= 1)
# nucleotide (O.C >= 0.5 & O.C < 1.7 & H.C > 1 & H.C < 1.8 & N.C >= 0.2 & N.C <= 0.5 & P.C >= 0.1 & P.C <= 0.35 &
N.P > 0.6 & N.P <= 5 & N >= 2 & P >= 1 & S == 0 & Mass > 305 & Mass < 523)
# However, there is a “gap space” between O.C > 0.6 & H.C >= 1.32 & H.C <= 1.45 for N =0 MF
```

```r
# First, make necessary elemental ratios
MF$O.C <- MF$O / MF$C # already present if using ICBM-OCEAN
MF$H.C <- MF$H / MF$C # already present if using ICBM-OCEAN
MF$N.C <- MF$N / MF$C
MF$P.C <- MF$P / MF$C
MF$N.P <- MF$N / MF$P # Many are 0 or Inf, not include in the rule
MF$DBE.C <- MF$DBE / MF$C
MF$DBE.H <- MF$DBE / MF$H
MF$DBE.O <- MF$DBE / MF$O

# Proposed integrated compound category classification (IC3) rule (without CRAM) -----------
# Laszakovits & Mackay (2022)'s MSCC is executed first as basis.
# Proteins 1&2 are denoted as peptide
# Fill the gap between O.C > 0.6 & H.C >= 1.32 & H.C <= 1.45 for N =0 MF by phytochemical and lipids, given possible allocations to this area (Figs S3&S5&S9 in Rivas-Ubach et al., 2018).
# lipid first, then phytochemical considering higher possibility of phytochemicals in this gap.
# This additional lipid allowed between O.C > 0.6 & O.C <= 0.8 & H.C >= 1.32 & H.C <= 1.8 (Fig. S3), but these lipids outside the gap space must be overwritten by aminosugar and carbohydrate for double classifications occur
MF$Category[MF$O.C <= 0.6 & MF$H.C >= 1.32 & MF$N.C <= 0.126 & MF$P.C < 0.35] <- "Lipid"
MF$Category[MF$O.C > 0.6 & MF$O.C <= 0.8 & MF$H.C >= 1.32 & MF$H.C <= 1.8 & MF$N.C <= 0.126 & MF$P.C < 0.35] <- "Lipid" # added
MF$Category[MF$O.C >= 0.56 & MF$H.C >= 1.53 & MF$H.C < 2.7 & MF$N == 0] <- "Carbohydrate"
MF$Category[MF$O.C >= 0.61 & MF$H.C >= 1.45 & MF$N.C <= 0.2 & MF$N.C > 0.07 & MF$P.C < 0.3 & MF$O >= 3 & MF$N >= 1] <- "Aminosugar"
MF$Category[MF$O.C <= 1.15 & MF$H.C < 1.32 & MF$N.C < 0.126 & MF$P.C <= 0.2] <- "Phytochemical"
MF$Category[MF$O.C > 0.6 & MF$O.C <= 1.15 & MF$H.C >= 1.32 & MF$H.C <= 1.45 & MF$N.C < 0.126 & MF$P.C <= 0.2] <- "Phytochemical" # added
MF$Category[MF$O.C > 0.12 & MF$O.C <= 0.6 & MF$H.C > 0.9 & MF$H.C < 2.5 & MF$N.C >= 0.126 & MF$N.C <= 0.7 & MF$P.C < 0.17 & MF$N >= 1] <- "Peptide"
MF$Category[MF$O.C > 0.6 & MF$O.C <= 1 & MF$H.C > 1.2 & MF$H.C < 2.5 & MF$N.C > 0.2 & MF$N.C <= 0.7 & MF$P.C < 0.17 & MF$N >= 1] <- "Peptide"
MF$Category[MF$O.C >= 0.5 & MF$O.C < 1.7 & MF$H.C > 1 & MF$H.C < 1.8 & MF$N.C >= 0.2 & MF$N.C <= 0.5 & MF$P.C >= 0.1 & MF$P.C <= 0.35 & MF$N >= 2 & MF$P >= 1 & MF$S == 0 & MF$mz > 304 & MF$mz < 522] <- "Nucleotide"

# Then, overwriting the above classification by the criteria in Laszakovits & Mackay (2022) for lignin and tannin,
# but with N = 0 to make their classifications more explicit and less influenced by the order of classifications.
# Additionally consider aromatic following the unequivocal criterion of Koch and Dittmar (2005/2015), overwriting all.
MF$Category[MF$O.C >= 0.21 & MF$O.C <= 0.44 & MF$H.C >= 0.86 & MF$H.C <= 1.34 & MF$N == 0] <- "Lignin"
MF$Category[MF$O.C >= 0.16 & MF$O.C <= 0.84 & MF$H.C >= 0.7 & MF$H.C <= 1.01 & MF$N == 0] <- "Tannin"
MF$Category[MF$AI.mod > 0.5] <- "Aromatic"
# Condensed aromatic can be further considered following Koch and Dittmar (2005/2015) if desired.
# MF$Category[MF$AI.mod >= 0.67] <- "CondensedAromatic"

# Check assignment results
dplyr::count(MF, Category)
# Nearly 100% assignment is possible
# Category    n
# 1     Aminosugar  117
# 2       Aromatic 1168
# 3   Carbohydrate  180
# 4         Lignin  491
# 5          Lipid 1968
# 6        Peptide  571
# 7 Phytochemical 1885
# 8         Tannin  413
# 9           <NA>   93
(6886-93)/6886*100 # 98.6% assignment
```

```
# Proposed integrated compound category classification (IC3) rule (with CRAM) -----------

# This is more suitable for NOM research when the presence of CRAM is predicted.
# Hertkorn et al. (2006) CRAM loosely defined by DBE/C (0.30–0.68), DBE/H (0.20–0.95), and DBE/O (0.77–1.75)

# Change the assignment order
# 1. Assign phytochemicals first to largely overwrite them by CRAM afterwards, as phytochemicals do not really make sense in NOM.
# Phytochemicals are not really needed because almost all of them will be overwritten by CRAM.
# 2. CRAM covers very wide space, so assign CRAM next and then overwrite it by other categories.
# This is to be conservative for CRAM assignment, as its assignment is not very constrained.
MF$Category[MF$O.C <= 1.15 & MF$H.C < 1.32 & MF$N.C < 0.126 & MF$P.C <= 0.2] <- "Phytochemical"
MF$Category[MF$O.C > 0.6 & MF$O.C <= 1.15 & MF$H.C < 1.45 & MF$N.C < 0.126 & MF$P.C <= 0.2] <- "Phytochemical" # added
MF$Category[MF$DBE.C >= 0.3 & MF$DBE.C <= 0.68] <- "CRAM"
MF$Category[MF$DBE.H >= 0.2 & MF$DBE.H <= 0.95] <- "CRAM"
MF$Category[MF$DBE.O >= 0.77 & MF$DBE.O <= 1.75] <- "CRAM"
MF$Category[MF$O.C <= 0.6 & MF$H.C >= 1.32 & MF$N.C <= 0.126 & MF$P.C < 0.35] <- "Lipid"
MF$Category[MF$O.C > 0.6 & MF$O.C <= 0.8 & MF$H.C >= 1.32 & MF$N.C <= 0.126 & MF$P.C < 0.35] <- "Lipid" # added
MF$Category[MF$O.C >= 0.56 & MF$H.C >= 1.53 & MF$H.C < 2.7 & MF$N == 0] <- "Carbohydrate"
MF$Category[MF$O.C >= 0.61 & MF$H.C >= 1.45 & MF$N.C <= 0.2 & MF$N.C > 0.07 & MF$P.C < 0.3 & MF$O >= 3 & MF$N >= 1] <- "Aminosugar"
MF$Category[MF$O.C > 0.12 & MF$O.C <= 0.6 & MF$H.C > 0.9 & MF$H.C < 2.5 & MF$N.C >= 0.126 & MF$N.C <= 0.7 & MF$P.C < 0.17 & MF$N >= 1] <- "Peptide"
MF$Category[MF$O.C > 0.6 & MF$O.C <= 1 & MF$H.C > 1.2 & MF$H.C < 2.5 & MF$N.C > 0.2 & MF$N.C <= 0.7 & MF$P.C < 0.17 & MF$N >= 1] <- "Peptide"
MF$Category[MF$O.C >= 0.5 & MF$O.C < 1.7 & MF$H.C > 1 & MF$H.C < 1.8 & MF$N.C >= 0.2 & MF$N.C <= 0.5 & MF$P.C >= 0.1 & MF$P.C <= 0.35 & MF$N >= 2 & MF$P >= 1 & MF$S == 0 & MF$mz > 304 & MF$mz < 522] <- "Nucleotide"
MF$Category[MF$O.C >= 0.21 & MF$O.C <= 0.44 & MF$H.C >= 0.86 & MF$H.C <= 1.34 & MF$N == 0] <- "Lignin"
MF$Category[MF$O.C >= 0.16 & MF$O.C <= 0.84 & MF$H.C >= 0.7 & MF$H.C <= 1.01 & MF$N == 0] <- "Tannin"
MF$Category[MF$AI.mod > 0.5] <- "Aromatic"

# Check assignment results
dplyr::count(MF, Category)
# Nearly 100% assignment is possible
# Category    n
# 1     Aminosugar  117
# 2        Aromatic 1168
# 3           CRAM 1866
# 4   Carbohydrate  180
# 5          Lignin  491
# 6           Lipid 2058
# 7         Peptide  571
# 8   Phytochemical    3
# 9          Tannin  413
# 10           <NA>   19
(6886-19)/6886*100 # 99.7% assignment, phytochemicals not necessary
# Un assigned MFs occur at the top-right corner of VK diagram, failing to be assigned as aminosugar or carbohydrate
# VK diagram without "Phytochemical" is shown in Fig. S1.
```

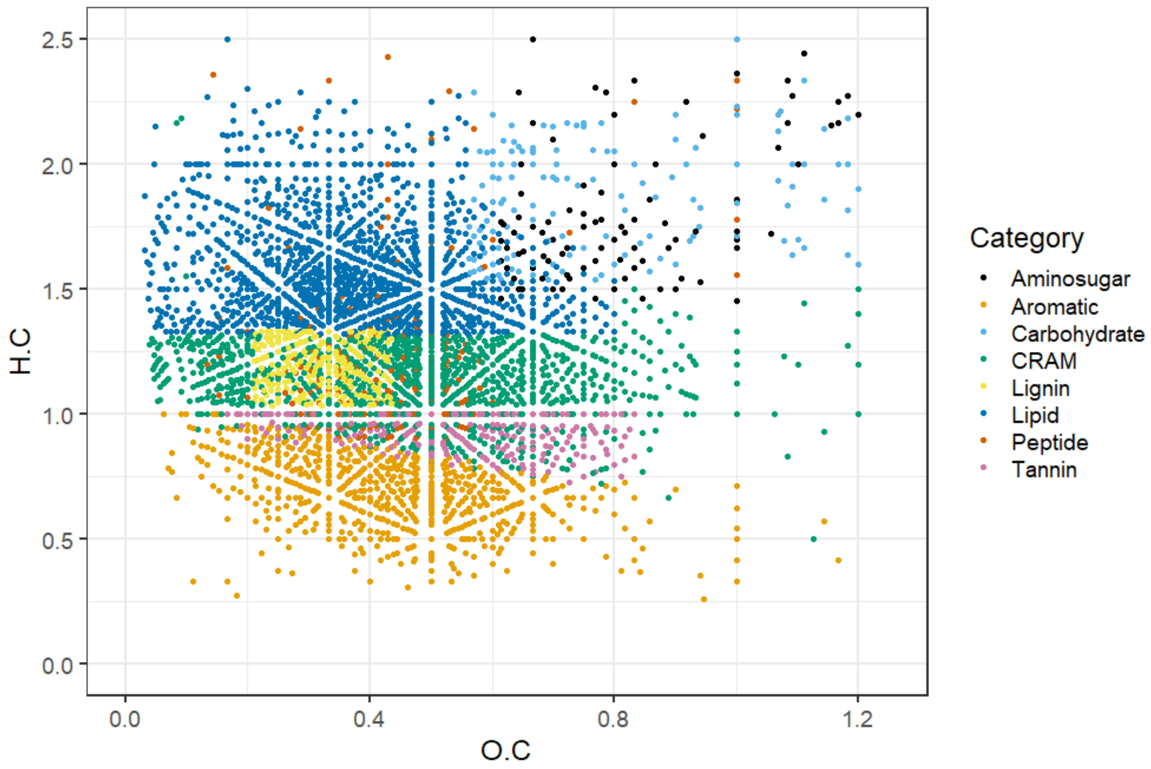


**Fig. S1.** The van Krevelen diagram and assigned compound categories by the proposed integrated compound category classification (IC3) rule.

**Supplementary Discussion S3**

**CCorA**

We evaluated the DOM-microbial linkage using two approaches: CCorA and network analysis. First, it was worth noting that different ASVs were detected to be co-varying with MFs depending on the method used, with differences even observed at the class level. In CCorA, the ASVs detected primarily belonged to *Planctomycetes* and *Verrucomicrobiae* (Fig. S4, Table S4), whereas network analysis predominantly identified *Actinobacteria*, *Anaerolineae*, *Gammaproteobacteria*, *Acidimicrobiia*, and *Bacteroidia* (Fig. 5, Table S5). These differences arose from method differences, and both approaches were considered valid. CCorA was conducted on the principal coordinates explaining the major (>75%) variance in the Bray-Curtis dissimilarity matrix of each SPE-DOM and microbial dataset. The correlated canonical axes thus comprised linear combinations of the principal coordinates, representing two specific dimensions in SPE-DOM and microbial space. CCorA so conducted captured a major canonical portion of the multivariate space between SPE-DOM and microbiome which may not be apparent in unconstrained ordination such as PCA (Anderson and Willis, 2003). In turn, post-hoc correlation of ASVs and MFs along the canonical axes may reflect covariation driven by a shared environmental factor rather than a direct relationship, as observed in an estuarine system where salinity serves as a co-factor (Osterholz et al., 2016). In our case, water mass mixing was unlikely to be a significant factor, given that sampling was conducted in a stratified lake with a pseudo-closed system. One of the advantages of CCorA is that it can be useful in interpreting the overall feature of the datasets by giving interpretable meaning in multivariate sample distributions, which is not directly possible using other multivariate correlation techniques, such as a co-occurrence network. In our CCorA results, CCorA1 captured processes occurring in the hypolimnion during water stratification (Fig. S3b). As shown in Fig. S4, red-colored MFs and ASVs decreased in relative abundance, whereas blue-colored ones increased in the hypolimnion during stratification. The increased CHO MFs were mostly Lipid or Lignin MFs, suggesting selective preservation of such refractory compounds in the aphotic hypolimnion. Contrarily, most of the increased CHON MFs were CRAM or Peptide MFs, likely produced by heterotrophic microbial activity (Hertkorn et al., 2006).

**Epilimnion network**

The two ASVs identified in the epilimnion network were both affiliated with order *Burkholderiales* within class *Gammaprotepbacteria* (Fig. 5a). Members of *Gammaprotepbacteria* are *r*-strategists that respond rapidly to the supply of labile organic matter, such as during phytoplankton blooms (Chiriac et al., 2023;

Francis et al., 2021). These bacteria are likely primary remineralizers of large polymeric organic matter, such as polysaccharides and proteins, which were out of our analytical window due to the constraints from PPL solid phase extraction (S1). However, the substrate-size preference of *Gammaprotepbacteria* appears to be less stringent than that of *Bacteroidetes*, another rapid responder to blooms, and these bacteria also use smaller compounds including amino acids, monosaccharides, dimethylsulfoniopropionate, terpenoids, and aromatics (Francis et al., 2021). Our network analysis showed that all considered compound categories covaried with *Burkholderiales*, with ASV_00026 (family *Comamonadaceae*) showing stronger associations with Lipid and Lignin MFs and ASV_00005 (family *Methylophilaceae*) with CRAM, Aromatic, and Peptide MFs (Fig. 5a). The causal relationship between ASVs and MFs cannot be resolved solely from the network analysis; however, background knowledge could help in understanding whether the compound is consumed as substrate or produced as metabolite. For instance, *Methylophilaceae* has been reported to preferentially use low molecular weight C1 compounds (Chiriac et al., 2023), thus its associations with CRAM, Aromatic, and Peptide MFs could be interpreted that these MFs may be metabolites of *Methylophilaceae* from the consumption of C1 compounds.

**Table S1. Basic water parameters, optical parameters, cell count, and SPE-DOC recovery of each sample.**

| Sample | Sampling date | Depth [m] | WT [°C] | EC25 [µS/cm] | Chl-Flu. [ppb] | Turbidity [FTU] | pH | DO [mg/L] | TOC [mgC L] | DOC [mgC/L] | POC [mgC/L] | $S_{275\_295}$ [$nm^{-1}$] | $SUVA_{254}$ [L/mgC/m] | Cell count [$10^6$/mL] | SPE-DOC recovery [%] |
|---|---|---|---|---|---|---|---|---|---|---|---|---|---|---|---|
| LB2204-17B-5m | 2022/04/14 | 5 | 11.2 | 114.3 | 1.78 | 0.63 | 8.31 | 13 | 1.317 | 1.014 | 0.303 | 0.024 | 1.41 | 3.22 | 48.1 |
| LB2204-17B-60m | 2022/04/14 | 60 | 7.6 | 117.3 | 0.45 | 0.36 | 7.14 | 10.8 | 1.086 | 0.909 | 0.177 | 0.0239 | 1.59 | 0.93 | 54.3 |
| LB2205-17B-5m | 2022/05/19 | 5 | 15.8 | 111.7 | 2.79 | 0.96 | 8.16 | 11.3 | 1.566 | 1.252 | 0.314 | 0.024 | 1.3 | 2.91 | 44.1 |
| LB2205-17B-TC | 2022/05/19 | 15 | 12.3 | 113.7 | 2.06 | 0.63 | 7.29 | 10.4 | 1.396 | 1.129 | 0.267 | 0.0247 | 1.34 | 2.27 | 44.6 |
| LB2205-17B-60m | 2022/05/19 | 60 | 7.9 | 116.9 | 0.47 | 0.32 | 7.04 | 10.6 | 1.032 | 0.94 | 0.092 | 0.0247 | 1.49 | 1.02 | 48.9 |
| LB2205-17B-85m | 2022/05/19 | 85 | 7.4 | 117.9 | 0.49 | 0.53 | 6.94 | 9.4 | 1.053 | 0.964 | 0.089 | 0.024 | 1.47 | 0.93 | 49.1 |
| LB2206-17B-5m | 2022/06/17 | 5 | 20.1 | 111.6 | 1.7 | 0.8 | 8.53 | 10.1 | 1.475 | 1.172 | 0.303 | 0.0259 | 1.35 | 3.93 | 45.1 |
| LB2206-17B-TC | 2022/06/17 | 17 | 11.8 | 113.9 | 1.38 | 0.48 | 7.54 | 9.3 | 1.308 | 1.052 | 0.256 | 0.0231 | 1.48 | 2.45 | 45.6 |
| LB2206-17B-60m | 2022/06/17 | 60 | 7.9 | 117.2 | 0.3 | 0.36 | 7.2 | 9.3 | 1.022 | 0.948 | 0.074 | 0.0227 | 1.56 | 1.1 | 47.8 |
| LB2206-17B-85m | 2022/06/17 | 85 | 7.6 | 117.8 | 0.41 | 0.52 | 7.12 | 8.8 | 1.042 | 0.973 | 0.069 | 0.023 | 1.49 | 1.26 | 47.1 |
| LB2207-17B-5m | 2022/07/21 | 5 | 26.5 | 110.2 | 0.87 | 0.52 | 8.08 | 8.5 | 1.428 | 1.162 | 0.266 | 0.0262 | 1.41 | 4.59 | 44.2 |
| LB2207-17B-TC | 2022/07/21 | 15 | 14.4 | 113.5 | 1.03 | 0.7 | 7.14 | 8 | 1.383 | 1.048 | 0.334 | 0.0233 | 1.58 | 2.6 | 46.7 |
| LB2207-17B-60m | 2022/07/21 | 60 | 7.8 | 117.3 | 0.28 | 0.22 | 7.06 | 9.4 | 0.966 | 0.911 | 0.056 | 0.0228 | 1.6 | 0.95 | 49.7 |
| LB2207-17B-85m | 2022/07/21 | 85 | 7.5 | 118.2 | 0.34 | 0.51 | 7 | 7.4 | 0.987 | 0.908 | 0.079 | 0.0224 | 1.65 | 1.21 | 50.6 |
| LB2208-17B-5m | 2022/09/02 | 5 | 27.7 | 108.8 | 1.04 | 0.57 | 7.69 | 8 | 1.488 | 1.159 | 0.33 | 0.0267 | 1.47 | 4.72 | 49.2 |
| LB2208-17B-TC | 2022/09/02 | 15 | 17.8 | 111.5 | 0.56 | 1.54 | 6.8 | 6.3 | 1.211 | 1.101 | 0.11 | 0.022 | 1.55 | 2.11 | 45.9 |
| LB2208-17B-60m | 2022/09/02 | 60 | 7.9 | 117.1 | 0.28 | 0.3 | 6.79 | 8.1 | 0.936 | 0.895 | 0.041 | 0.0235 | 1.58 | 0.94 | 50 |
| LB2208-17B-85m | 2022/09/02 | 85 | 7.6 | 118.2 | 0.43 | 2.29 | 6.62 | 4.3 | 1.041 | 0.934 | 0.107 | 0.0228 | 1.57 | 1.21 | 51.4 |
| LB2209-17B-5m | 2022/09/22 | 5 | 24.1 | 109.2 | 1.03 | 0.59 | 6.95 | 7.8 | 1.406 | 1.194 | 0.212 | 0.0253 | 1.44 | 3.37 | 47.1 |
| LB2209-17B-TC | 2022/09/22 | 19 | 13.2 | 113.1 | 0.63 | 1.45 | 6.58 | 6.9 | 1.466 | 1.192 | 0.274 | 0.0251 | 1.44 | 2.94 | 46.6 |
| LB2209-17B-60m | 2022/09/22 | 60 | 7.9 | 117.4 | 0.3 | 0.89 | 6.47 | 7 | 0.993 | 0.918 | 0.075 | 0.0239 | 1.55 | 0.97 | 56.6 |
| LB2209-17B-85m | 2022/09/22 | 85 | 7.6 | 118.4 | 0.41 | 2.14 | 6.35 | 3.5 | 1.082 | 0.965 | 0.117 | 0.0222 | 1.6 | 0.98 | 56.6 |
| LB2210-17B-5m | 2022/10/28 | 5 | 19 | 110.5 | 1.7 | 0.49 | 7.1 | 9.3 | 1.497 | 1.221 | 0.276 | 0.0249 | 1.43 | 2.13 | 51.2 |
| LB2210-17B-TC | 2022/10/28 | 21 | 12.5 | 114 | 0.82 | 0.38 | 6.67 | 6.8 | 1.384 | 1.163 | 0.221 | 0.0245 | 1.42 | 2.08 | 55.2 |
| LB2210-17B-60m | 2022/10/28 | 60 | 7.9 | 117.3 | 0.26 | 0.28 | 6.34 | 7.3 | 0.941 | 0.933 | 0.008 | 0.0229 | 1.49 | 0.9 | 58.3 |
| LB2210-17B-85m | 2022/10/28 | 85 | 7.6 | 119.8 | 0.33 | 1.35 | 6.1 | 2.4 | 1.052 | 1.001 | 0.051 | 0.0216 | 1.6 | 1.14 | 58.7 |
| LB2211-17B-5m | 2022/11/18 | 5 | 16.5 | 111.4 | 1.32 | 0.46 | 6.89 | 9.4 | 1.457 | 1.18 | 0.277 | 0.0247 | 1.46 | 2.08 | 59.6 |
| LB2211-17B-TC | 2022/11/18 | 26 | 11.7 | 114.2 | 0.38 | 0.43 | 6.56 | 7.2 | 1.294 | 1.111 | 0.183 | 0.0241 | 1.46 | 2.19 | 56.5 |
| LB2211-17B-60m | 2022/11/18 | 60 | 8.1 | 116.9 | 0.29 | 0.44 | 6.09 | 6.4 | 1.049 | 0.964 | 0.085 | 0.0229 | 1.54 | 0.85 | 56.2 |
| LB2211-17B-85m | 2022/11/18 | 85 | 7.7 | 118.3 | 0.3 | 0.77 | 6.06 | 5.6 | 1.016 | 0.935 | 0.081 | 0.0214 | 1.67 | 0.89 | 61 |
| LB2212-17B-5m | 2022/12/22 | 5 | 11.2 | 114.2 | 1.38 | 1.01 | 7.17 | 10.3 | 1.331 | 1.159 | 0.172 | 0.0248 | 1.41 | 1.79 | 48.5 |
| LB2212-17B-TC | 2022/12/22 | 47 | 8.9 | 115.8 | 0.32 | 0.45 | 6.83 | 7.1 | 0.979 | 0.944 | 0.035 | 0.0254 | 1.44 | 0.99 | 50.3 |
| LB2212-17B-60m | 2022/12/22 | 60 | 8.2 | 116.6 | 0.3 | 0.29 | 6.57 | 7.2 | 0.945 | 0.929 | 0.015 | 0.0233 | 1.49 | 0.9 | 53.8 |
| LB2212-17B-85m | 2022/12/22 | 85 | 7.7 | 120.7 | 0.56 | 1.36 | 6.33 | 2.5 | 1.038 | 0.981 | 0.057 | 0.0186 | 1.69 | 1.03 | 51.9 |

WT = water temperatute, EC25 = electrical conductivity at 25℃, Chl-Flu = Chlorophyll fluorescence, DO = dissolved oxygen, TOC = total organic carbon, DOC = dissolved organic carbon <0.1 µm in size, POC = particulate organic carbon 250-0.1 µm in size, $S_{275-295}$ = spectral slope between 275-295 nm, $SUVA_{254}$ = DOC-specific UV sbsorbance at 254 nm, SPE-DOC recovery = DOC extraction efficiency by PPL resin.

**Table S2. Comparison of intensity-weighted mean and standard deviation (SD) for four metrics (oxygenation, hydrogen saturation, mass to charge ratio, and modified aromaticity index) and four compound classes of Suwannee River natural organic matter (SRNOM; 2R101N) in the negative-ion mode between this study and the reference values (Hawkes et al., 2020).**

| | This study (n = 6) | | Reference | |
|---|---|---|---|---|
| Parameters | Mean | SD | Mean | SD |
| H/C | 1.04 | 0.0 | 1.05 | 0.016 |
| O/C | 0.58 | 0.0 | 0.57 | 0.022 |
| m/z | 379 | 6.8 | 405 | 17 |
| $AI_{mod}$ | 0.35 | 0.0 | 0.34 | 0.009 |
| High O unsaturated | 60 | 1.1 | 63 | 5.8 |
| Low O unsaturtated | 25 | 1.4 | 26 | 5.5 |
| Aromatics | 14 | 1.0 | 10 | 1.6 |
| Aliphatic | 1.2 | 0.3 | 0.21 | 0.078 |

High O unsaturated = formulas with H/C < 1.5 and $AI_{mod}$ < 0.5 and O/C >= 0.5, Low O unsaturated = formulas with H/C < 1.5 and $AI_{mod}$ < 0.5 and O/C < 0.5, Aromatic = formulas with $AI_{mod}$ ≥ 0.5, Aliphatic = formulas with H/C ≥ 1.5.

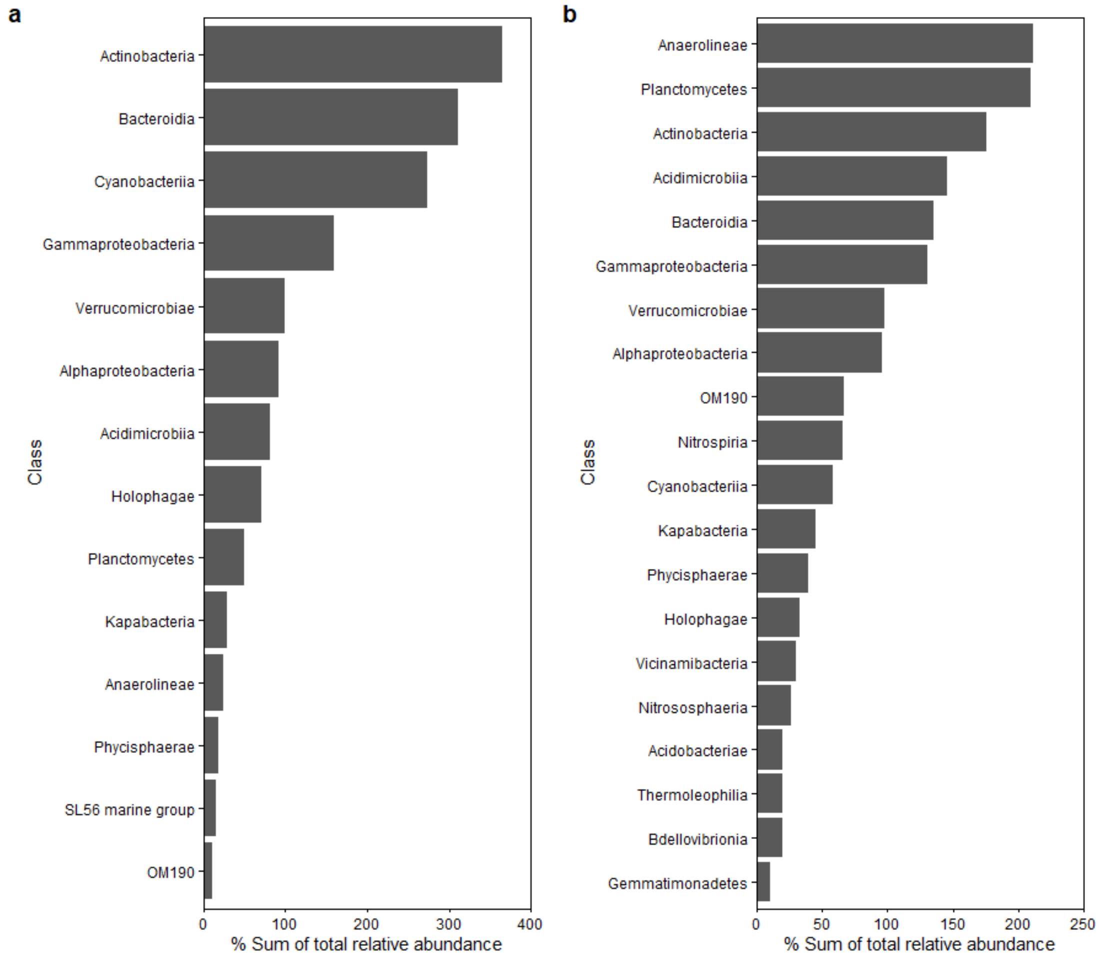


**Fig. S2. A class-resolved microbial community composition for epilimnion (5 m and TC) (a) and hypolimnion (60 m and 85 m) (b).** Only microbial classes of which sum of relative abundance across datasets higher than 10% are presented.

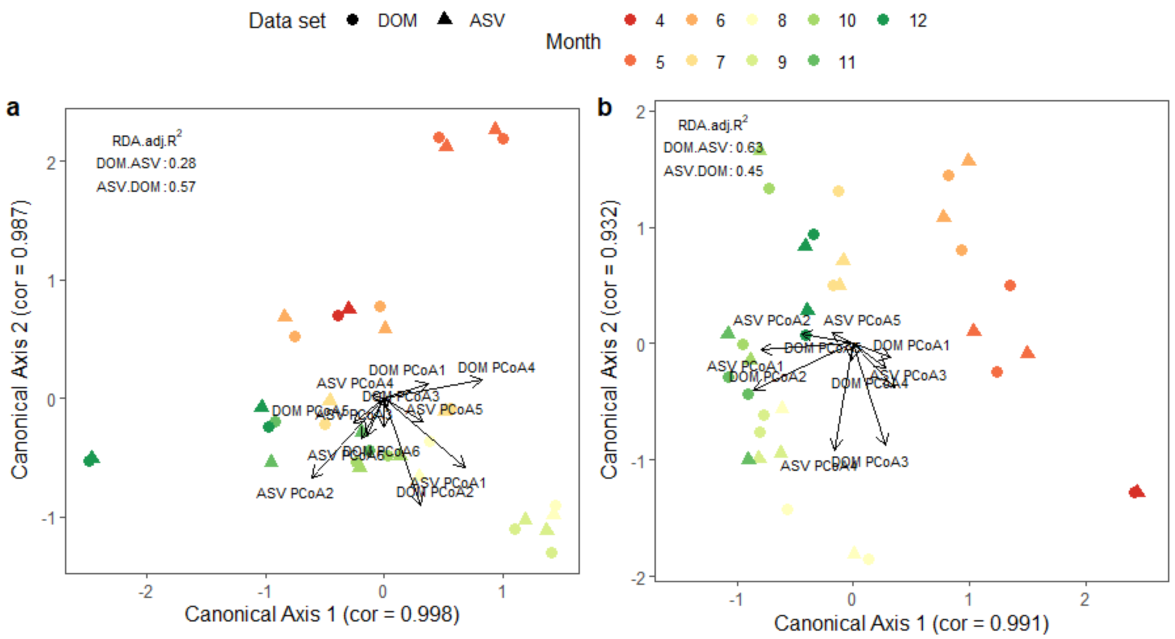


**Fig. S3. Canonical correlation analysis between SPE-DOM and microbiome for epilimnion (5 m and TC) (a) and hypolimnion (60 m and 85 m) (b).** DOM.ASV and ASV.DOM indicates explanatory powers of DOM on ASV and vice versa, respectively, calculated by bimultivariate redundancy coefficients (RDA.adj.$R^2$).

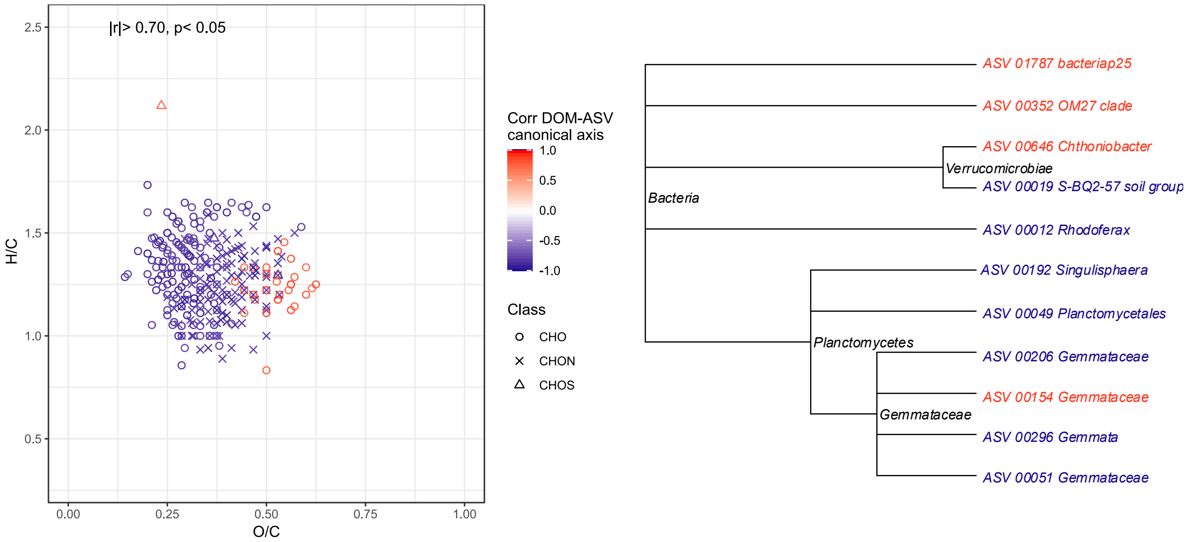


**Fig. S4. Co-varying molecular formulae and ASVs in the hypolimnion during lake water stratification.** Co-variation was assessed by the common significant correlations with the first canonical axis in canonical correlation analysis. Molecular formulae and ASVs in red decreased, while those in blue increased, in relative abundance during stratification in Lake Biwa. In the phylogenetic cluster, the label names are the combination of ASV ID and the lowest identifiable phylogenetic level (order to genus level).

**Table S3. Detailed taxonomic information of the hypolimnetic ASVs identified in canonical correlation analysis as having significant correlations with the first canonical axis.**

| ASV_ID | Kingdom | Phylum | Class | Order | Family | LabelName |
|---|---|---|---|---|---|---|
| ASV_00352 | Bacteria | Bdellovibrionota | Bdellovibrionia | Bdellovibrionales | Bdellovibrionaceae | ASV_00352_OM27 clade |
| ASV_01787 | Bacteria | Myxococcota | bacteriap25 | ND | ND | ASV_01787_bacteriap25 |
| ASV_00051 | Bacteria | Planctomycetota | Planctomycetes | Gemmatales | Gemmataceae | ASV_00051_Gemmataceae |
| ASV_00296 | Bacteria | Planctomycetota | Planctomycetes | Gemmatales | Gemmataceae | ASV_00296_Gemmata |
| ASV_00154 | Bacteria | Planctomycetota | Planctomycetes | Gemmatales | Gemmataceae | ASV_00154_Gemmataceae |
| ASV_00206 | Bacteria | Planctomycetota | Planctomycetes | Gemmatales | Gemmataceae | ASV_00206_Gemmataceae |
| ASV_00192 | Bacteria | Planctomycetota | Planctomycetes | Isosphaerales | Isosphaeraceae | ASV_00192_Singulisphaera |
| ASV_00049 | Bacteria | Planctomycetota | Planctomycetes | Planctomycetales | ND | ASV_00049_Planctomycetales |
| ASV_00012 | Bacteria | Proteobacteria | Gammaproteobacteria | Burkholderiales | Comamonadaceae | ASV_00012_Rhodoferax |
| ASV_00646 | Bacteria | Verrucomicrobiota | Verrucomicrobiae | Chthoniobacterales | Chthoniobacteraceae | ASV_00646_Chthoniobacter |
| ASV_00019 | Bacteria | Verrucomicrobiota | Verrucomicrobiae | S-BQ2-57 soil group | ND | ASV_00019_S-BQ2-57 soil group |

**Table S4. Detailed taxonomic information of the ASVs identified in the network analysis.** ASVs in green are all from family Sporichthyaceae (known as acI clade) showing tight links in the network, while ASVs in blue are all from genus CL500-29 marine group (known as acIV clade) but showing no links in the network. ASVs in purple are those identified as the hypolimnion specialists in past studies, while ASVs in orange are those found in the epilimnion network.

| ASV_ID | Kingdom | Phylum | Class | Order | Family | Genus |
|---|---|---|---|---|---|---|
| ASV_00001 | Bacteria | Actinobacteriota | Actinobacteria | Frankiales | Sporichthyaceae | hgcI clade |
| ASV_00003 | Bacteria | Chloroflexi | Anaerolineae | Anaerolineales | Anaerolineaceae | NA |
| ASV_00005 | Bacteria | Proteobacteria | Gammaproteobacteria | Burkholderiales | Methylophilaceae | Candidatus Methylopumilus |
| ASV_00006 | Bacteria | Actinobacteriota | Actinobacteria | Frankiales | Sporichthyaceae | Candidatus Planktophila |
| ASV_00009 | Bacteria | Actinobacteriota | Acidimicrobiia | Microtrichales | Ilumatobacteraceae | CL500-29 marine group |
| ASV_00014 | Bacteria | Planctomycetota | Phycisphaerae | Phycisphaerales | Phycisphaeraceae | CL500-3 |
| ASV_00015 | Bacteria | Actinobacteriota | Actinobacteria | Frankiales | Sporichthyaceae | hgcI clade |
| ASV_00022 | Bacteria | Actinobacteriota | Actinobacteria | Frankiales | Sporichthyaceae | hgcI clade |
| ASV_00026 | Bacteria | Proteobacteria | Gammaproteobacteria | Burkholderiales | Comamonadaceae | Acidovorax |
| ASV_00037 | Bacteria | Actinobacteriota | Actinobacteria | Frankiales | Sporichthyaceae | hgcI clade |
| ASV_00048 | Bacteria | Actinobacteriota | Acidimicrobiia | Microtrichales | Ilumatobacteraceae | CL500-29 marine group |
| ASV_00069 | Bacteria | Actinobacteriota | Actinobacteria | Frankiales | Sporichthyaceae | Candidatus Planktophila |
| ASV_00070 | Bacteria | Acidobacteriota | Vicinamibacteria | Vicinamibacterales | Vicinamibacteraceae | NA |
| ASV_00083 | Bacteria | Proteobacteria | Alphaproteobacteria | Reyranellales | Reyranellaceae | Reyranella |
| ASV_00131 | Bacteria | Acidobacteriota | Acidobacteriae | Paludibaculum | NA | NA |
| ASV_00156 | Bacteria | Actinobacteriota | Acidimicrobiia | Microtrichales | Ilumatobacteraceae | CL500-29 marine group |
| ASV_00159 | Bacteria | Proteobacteria | Gammaproteobacteria | Burkholderiales | NA | NA |
| ASV_00162 | Bacteria | Cyanobacteria | Cyanobacteriia | Chloroplast | NA | NA |
| ASV_00268 | Bacteria | Bacteroidota | Bacteroidia | Chitinophagales | Chitinophagaceae | Sediminibacterium |
| ASV_00341 | Bacteria | Bacteroidota | Kapabacteria | Kapabacteriales | NA | NA |

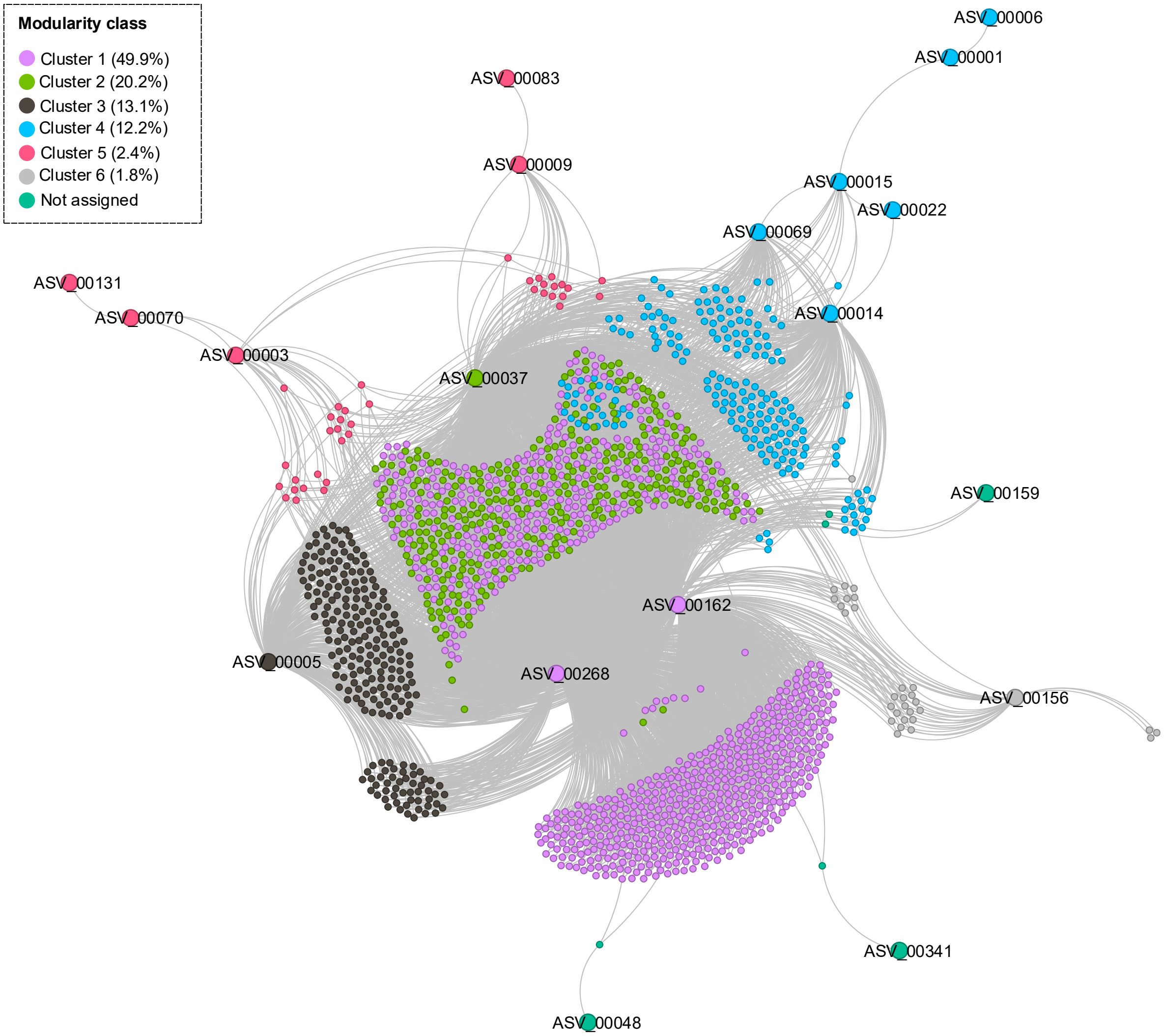


**Fig. S5. Network modularity class of molecular formulae and ASVs in the hypolimnion.** Molecular formulae and ASVs are color-coded by modularity class. Numbers in parentheses atter the modularity class numbers indicate the percentages of the numbers of nodes in each cluster.

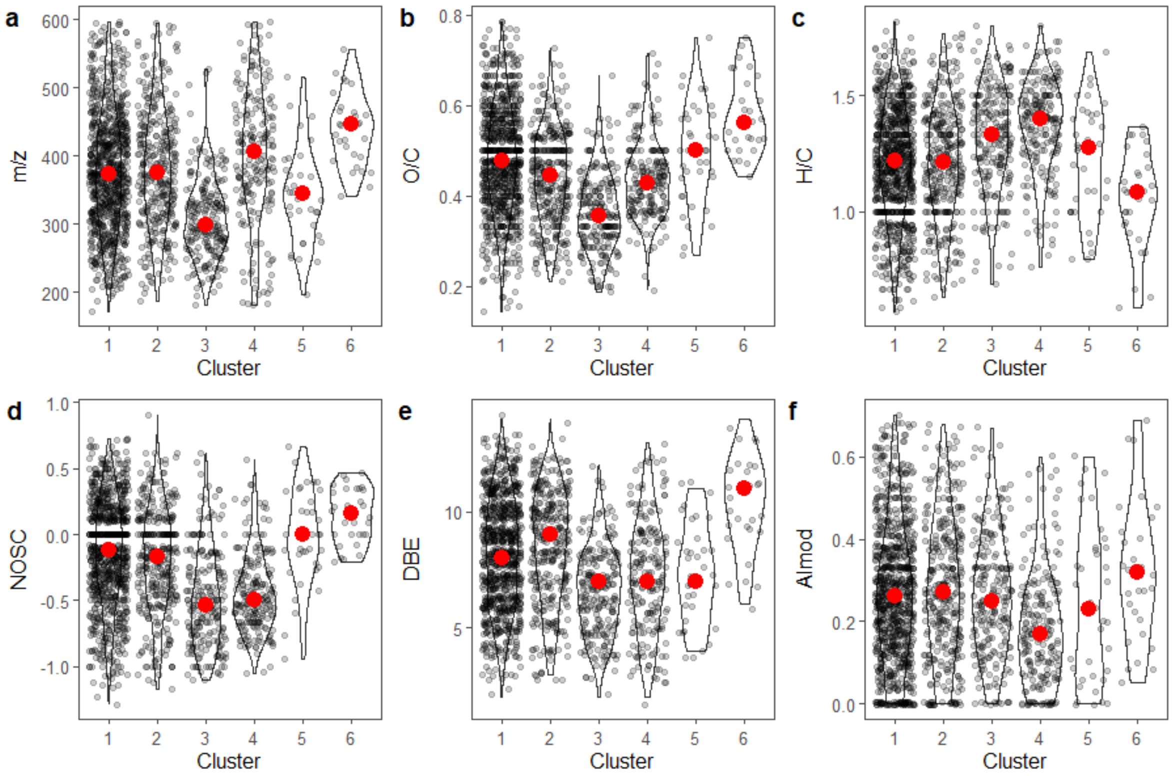


**Fig. S6. Molecular properties of molecular formulae in each cluster identified in Fig. S4.** Black dots represent each sample, violin plots are sample distributions, and the red points indicate the medians.

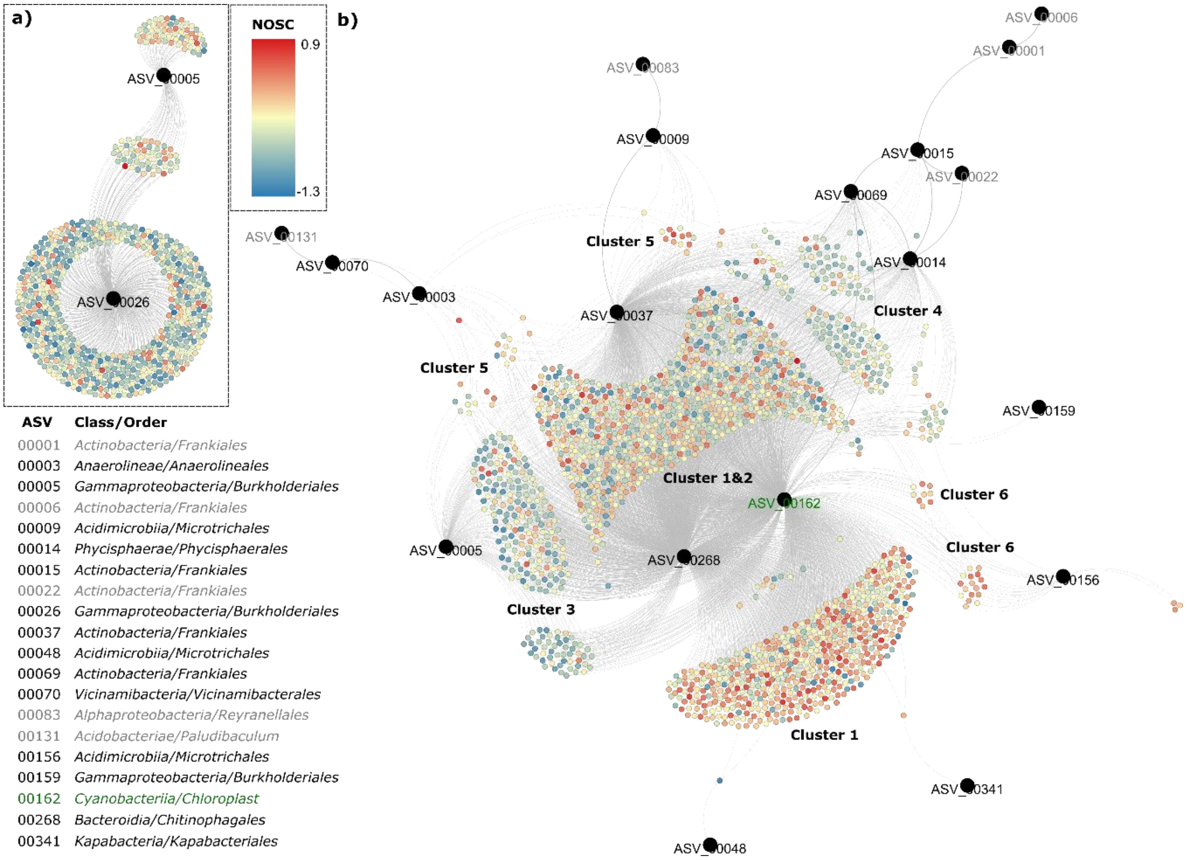


**Fig. S7. Network of molecular formulae and ASVs in (a) the epilimnion and (b) the hypolimnion during lake water stratification.** The positive network was constructed by the proportionality index of parts (PIP). Molecular formulae are color-coded by nominal oxidation state of carbon (NOSC), while ASVs are shown in larger black circles. The class and order of each ASV are presented after the corresponding ASV numbers. Grey ASVs are those without direct linkage with any molecular formula, and a green ASV is a chloroplast (see main manuscript for discussion).
